# Estimating correlated rates of trait evolution with uncertainty

**DOI:** 10.1101/102939

**Authors:** D.S. Caetano, L.J. Harmon

**Affiliations:** Department of Biological Sciences, Institute for Bioinformatics and Evolutionary Studies (IBEST), University of Idaho, Moscow, Idaho, 83843, USAs

**Keywords:** Anolis, Centrarchidae, comparative methods, evolutionary integration, evolutionary rates, modularity, pruning algorithm

## Abstract

Correlated evolution among traits, which can happen due to genetic constraints, ontogeny, and selection, can have an important impact on the trajectory of phenotypic evolution. For example, shifts in the pattern of evolutionary integration may allow the exploration of novel regions of the morphospace by lineages. Here we use phylogenetic trees to study the pace of evolution of several traits and their pattern of evolutionary correlation across clades and over time. We use regimes mapped to the branches of the phylogeny to test for shifts in evolutionary integration. Our approach incorporates the uncertainty related to phylogeny, ancestral state estimates and parameter estimates to produce posterior distributions using Bayesian Markov chain Monte Carlo. We implemented the use of summary statistics to test for regime shifts based on a series of attributes of the model that can be directly relevant to biological hypotheses. In addition, we extend Felsenstein’s pruning algorithm to the case of multivariate Brownian motion models with multiple rate regimes. We performed extensive simulations to explore the performance of the method under a series of scenarios. Finally, we provide two test cases; the evolution of a novel buccal morphology in fishes of the family Centrarchidae and a shift in the trajectory of evolution of traits during the radiation of anole lizards to and from the Caribbean islands.

---

Correlated evolution among traits, known as evolutionary integration, is ubiquitous across the tree of life and can have an important impact on the trajectory of phenotypic evolution (Olson and Miller, 1958; Klingenberg and Marugán-Lobón, 2013; Armbruster et al., 2014; Klingenberg, 2014; Goswami et al., 2014, 2015; Melo et al., 2016). Genetic constraints, ontogeny, and selection have pivotal roles in the development and maintenance of morphological integraion over time (Arnold, 1992; Arnold et al., 2001; Hansen and Houle, 2004; Goswami et al., 2015; Melo et al., 2016). When the additive genetic covariance between traits is strong, then evolutionary correlation is likely due to genetic factors. In contrast, traits might not show strong genetic covariance and still be evolutionarily integrated due to correlated selection, which can be a result of distinct factors, such as anatomical interactions during growth or coordinated function (Armbruster and Schwaegerle, 1996; Armbruster et al., 2014). For instance, correlated evolution can be favored by selection to maintain a cohesive pattern of variation among traits with a shared function, but evolution can be hindered if the genetic covariance is not aligned with the selection gradient (Lande, 1979; Schluter, 1996; Villmoare, 2012; Goswami et al., 2014). Alternatively, when evolutionary correlation is mainly a result of correlated selection then the morphospace occupied by lineages can be restricted by the strength and direction of the selection gradient (Felsenstein, 1988; Armbruster and Schwaegerle, 1996). Shifts in the pattern of evolutionary integration among traits over macroevolutionary scales, due to changes in the genetic architecture or selection gradient, may play a fundamental role in the exploration of novel regions of the morphospace (Young and Hallgr´ımsson, 2005; Goswami, 2006; Revell and Collar, 2009; Monteiro and Nogueira, 2010; Hallgr´ımsson et al., 2012; Claverie and Patek, 2013).

Macroevolutionary transitions in morphospace evolution have been associated with both increases and decreases in the evolutionary integration among traits. In centrarchid fishes, for example, the evolution of a novel mouth morphology was followed by a rapid differentiation of feeding habits (Collar et al., 2005). More specifically, the increase in the evolutionary correlation between two morphological features of the suction-feeding mechanism in species of *Micropterus* is associated with a specialization towards consumption of larger prey (Collar et al., 2005; Revell and Collar, 2009). In contrast, the once strong developmental integration between the fore- and hindlimbs of early tetrapods underwent a dramatic change allowing the limbs to respond to diverging selective pressures and leading to the evolution of bipedalism and flight (Young and Hallgr´ımsson, 2005; Young et al., 2010; Dececchi and Larsson, 2013). These examples show the role of shifts in evolutionary integration associated with the evolution of novel morphologies. However, stable patterns of evolutionary integration over long time scales can be responsible for the constraint of lineages to limited regions of the morphospace and might be a plausible mechanism associated with patterns of stasis observed in the fossil record (Hansen and Houle, 2004; Bolstad et al., 2014; Goswami et al., 2015). Thus, evolutionary trait correlations are central to the maintenance of form and function through time and can either drive or slow morphological differentiation.

Despite the prevalent role of evolutionary integration, most of what we know about the tempo and mode of trait evolution come from studies of single traits (e.g., Harmon et al., 2010; Hunt et al., 2015, among others). Even when multiple traits are the object of investigation, studies often use principal component axes (or phylogenetic PCA; Revell, 2009) to reduce the dimensionality of the data so that univariate methods can be applied (Harmon et al., 2010; Mahler et al., 2013; Klingenberg and Marug´an-Lob´on, 2013, see Uyeda et al. 2015 for more examples). This is most likely a reflection of the phylogenetic comparative models of trait evolution available for use, since few are focused on two or more traits (but see Revell and Harmon, 2008; Hohenlohe and Arnold, 2008; Revell and Collar, 2009; Bartoszek et al., 2012; Adams, 2012, 2014b; Clavel et al., 2015; Adams and Collyer, 2017; Bastide et al., 2018). However, studying one trait at a time eliminates the possibility of identifying patterns of evolutionary correlation, while principal component axes does not allow testing for evolutionary shifts in integration because the orientation of the PC axes are homogeneous across the branches of the phylogenetic tree. Furthermore, it also has been shown that PCA can influence our biological interpretation about the mode of evolution of the data (Uyeda et al., 2015) because the first PC axes are consistently estimated as early bursts of differentiation whereas the last axes store a strong signal of stabilizing selection, independent of the true model of evolution of the traits. As a result, we need models that apply to multivariate data as such in order to better understand macroevolutionary patterns of evolutionary integration.

One way to model multivariate trait evolution using phylogenetic trees is through the evolutionary rate matrix (Hohenlohe and Arnold, 2008; Revell and Harmon, 2008; Revell and Collar, 2009; Adams and Felice, 2014). This is a variance-covariance matrix that describes the rates of trait evolution under Brownian motion in the diagonals and the evolutionary covariance among traits (i.e., the pattern of evolutionary integration) in the off-diagonals (Huelsenbeck and Rannala, 2003; Revell and Harmon, 2008). The evolutionary rate matrix is ideal for studying patterns of evolutionary integration because it allows for simultaneous estimate of the individual rates of evolution of each trait as well as the evolutionary covariance between each pair of traits. It is also a flexible model, since any number of evolutionary rate matrix regimes can be fitted to the same phylogenetic tree (Revell and Collar, 2009). The contrast between evolutionary rate matrices independently estimated in different regions of the tree can inform us about the magnitude and direction of shifts in the pattern of evolutionary integration.

One of the challenges of working with rate matrices is that covariances can be hard to estimate, especially when the number of species (observations) is small relative to the number of traits (parameters) in the model (Adams and Collyer, 2017). As the number of parameters in a model increases, the amount of data required for proper estimation also increases and it becomes crucial to directly incorporate uncertainty in parameter estimates when interpreting results. However, the majority of studies to date have relied on point estimates of the evolutionary rate matrix by maximum likelihood (Revell and Harmon, 2008; Revell and Collar, 2009; Clavel et al., 2015; Goolsby, 2016, but see Huelsenbeck and Rannala, 2003 and Dines et al., 2014 for exceptions). Although the confidence interval around the maximum likelihood estimate can be used as a measure of uncertainty, this quantity is rarely reported (Revell and Harmon, 2008; Revell and Collar, 2009; Adams, 2012; Immler et al., 2012; Adams and Felice, 2014; Collar et al., 2014). Furthermore, the uncertainty in parameter estimates does not take direct part in model selection using likelihood ratio tests or AIC (Burnham and Anderson, 2003), which can lead researchers to erroneous conclusions about their models (Boettiger et al., 2012). Besides the possible uncertainty in parameter estimates, there is an important computational burden associated with the evaluation of the likelihood function of the multivariate Brownian motion model due to the computation of matrix inversions and determinants (Felsenstein, 1973; Hadfield and Nakagawa, 2010; Freckleton, 2012). Thus, computational time can become a limitation when performing a large number of likelihood evaluations, such as in simulation based approaches.

Recently, Adams (2014b) described a method to estimate the rate of evolution under Brownian motion of traits defined by several dimensions (high-dimensional data), even when the number of trait dimensions exceeds the number of lineages in the phylogeny. This method was extended to a plethora of variations based on the same general framework (Adams and Felice, 2014; Adams, 2014a; Denton and Adams, 2015; Adams and Collyer, 2017, see also Goolsby (2016) for a different implementation). These methods work with high-dimensional data as a result of the use of distance matrices rather than covariance matrices, since the later becomes singular if the number of variables (traits) is larger than the number of observations (species). However, by avoiding the calculation of the covariance among trait dimensions (Adams, 2014b), such suite of methods assume a homogeneous rate of evolution and a common structure of evolutionary correlation shared by all dimensions of the trait. Thus, 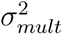 is ideal for high-dimensional traits such as shape data (Adams, 2014b), but it has important limitations for the study of evolutionary integration among multiple traits, which is the main focus of this study.

In order to ask questions about the evolution of integration using phylogenetic trees we need a computationally efficient method that can estimate evolutionary rate matrices while incorporating uncertainty in parameter estimates. Here we implement a Bayesian estimate for the evolutionary rate matrix using Markov chain Monte Carlo (MCMC) to provide a direct assessment of the uncertainty associated with parameter estimates in the form of a posterior distribution. Our implementation also allows for multiple regime configurations and/or phylogenetic trees to be incorporated in the MCMC chain, thus integrating the uncertainty associated with ancestral state estimates and phylogenetic reconstruction to the analysis. In order to increase the performance of the likelihood evaluation, we implemented Felsenstein’s (1973) pruning algorithm. For this, we derive a new version of the pruning algorithm that is suitable for the special case when several rate regimes of the multivariate Brownian motion model are fit to different branches of the same phylogenetic tree. We apply our new approach to two biological examples: the fast evolution of morphology associated with the radiation of *Anolis* lizards from mainland South America to the Caribbean islands (Pinto et al., 2008; Mahler et al., 2013; Moreno-Arias and Calder´on-Espinosa, 2016; Poe et al., 2018) and the shift of feeding habits driven by the change in mouth morphology in Centrarchidae fishes (Revell and Collar, 2009). We show that there is no detectable shift in the evolutionary integration among morphological traits during the anole radiation and that, given the significant uncertainty in estimates of evolutionary correlation, it is unlikely that a shift in the pattern of integration happened in Centrarchidae. We also provide results from extensive simulations showing that our approach has good performance under diverse scenarios of correlated evolution.

## Methods

Our new approach implements a multivariate Brownian motion model with a single or multiple rate regimes fitted to the phylogenetic tree. The number and position of rate regimes mapped on the branches of the phylogeny can be informed by a discrete trait distributed among the tips of the tree. For this, we reconstruct the ancestral state of the predictive trait at the branches of the tree and associate a different rate regime for each state (Huelsenbeck et al., 2003; Revell, 2012, see Fig. 1). Then we use MCMC to simultaneously estimate the posterior distribution for the rates of evolution of each continuous trait as well as their evolutionary covariation in the form of an evolutionary rate matrix (**R**; Revell and Harmon, 2008; Revell and Collar, 2009). Our primary objective is to provide a framework to incorporate uncertainty in the estimates of **R** as well as to build a flexible model to study shifts in evolutionary correlation across clades and over time. Our method requires a phylogenetic tree with branch lengths, continuous data for two or more traits for each tip species, and it uses Metropolis-Hastings Markov chain Monte Carlo (MCMC, Metropolis et al., 1953; Hastings, 1970).

**Figure 1:**
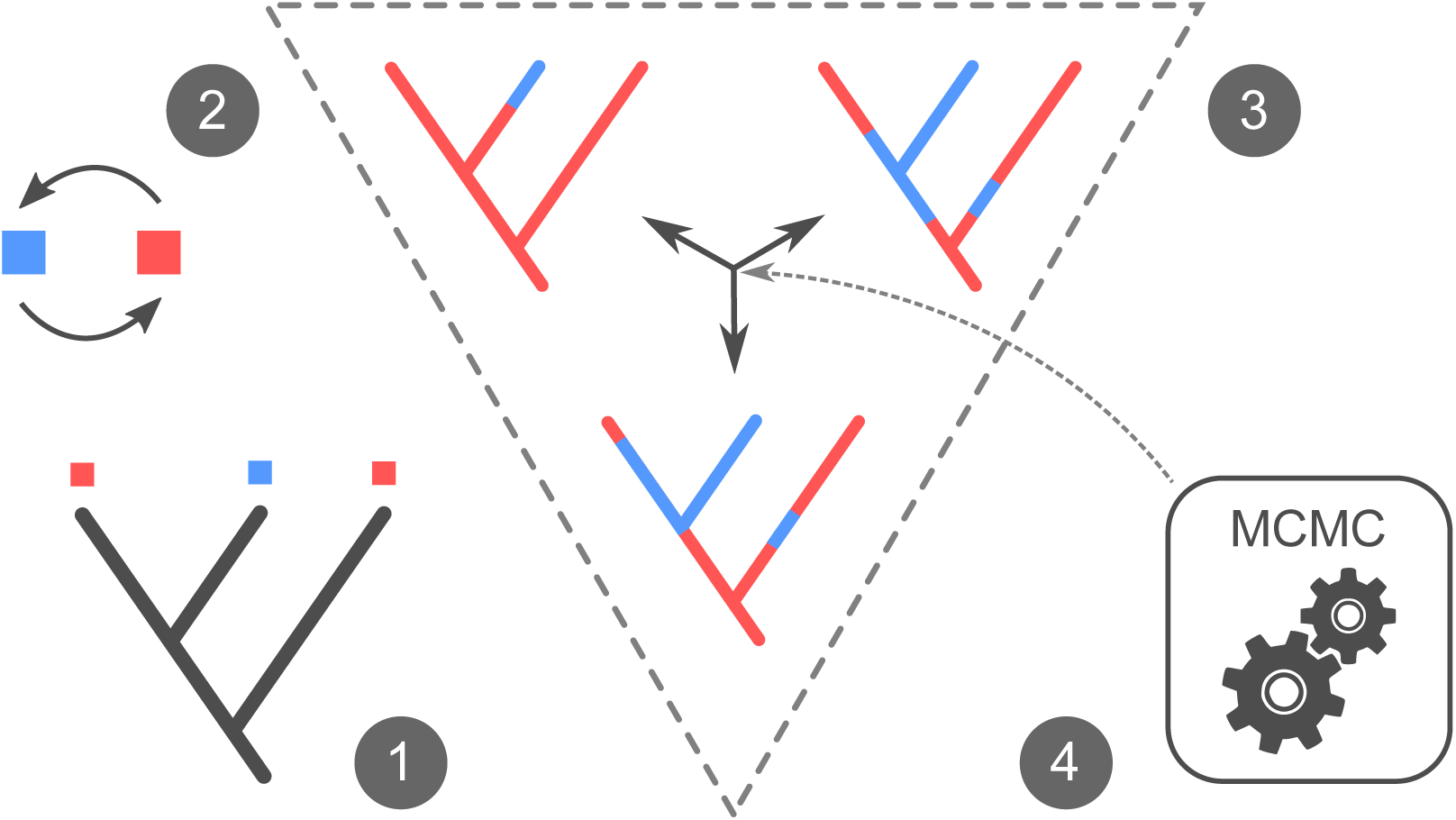
Example of workflow for our approach to incorporate uncertainty in ancestral state reconstruction and regime configurations into the posterior distribution of parameter estimates. Observed states at the tips of the phylogeny for the predictive categorical trait (1) are used to estimate the transition rate matrix (2). A collection of stochastic character histories is simulated (3) based on the rates of transition between the states (2). The Markov chain Monte Carlo (MCMC) computes the likelihood of the model given a rate regime configuration. At every step of the MCMC a new stochastic character history is sampled from the pool (4). As a result, the posterior distribution of parameter estimates for the model incorporates the uncertainty associated with ancestral state reconstructions (3).

### The model

The evolutionary rate matrix (**R**; Revell and Harmon, 2008; Revell and Collar, 2009) is a variance-covariance matrix with rates of evolution for each trait in the diagonal and the evolutionary covariance between each pair of traits in the off-diagonals. Revell and Collar (2009) derived a general form of the likelihood function for the model that allows for several independent matrices assigned to different branches of the phylogenetic tree

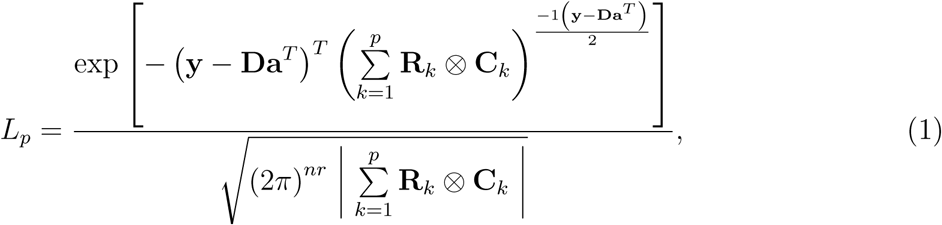

where **y** is a vector of length *n ⋅ r* derived by concatenating the columns of a *n* by *r* matrix of trait values for *n* tips and *r* traits; **D** is a *n ⋅ r* by *r* design matrix composed of 1 for each (*i, j*) entry that satisfies (*j −* 1) *⋅ n < i ≤ j ⋅ n* and 0 otherwise; **a** is a vector with *r* root values for the tree (or the phylogenetic mean); **R**_*k*_is the *k*th evolutionary rate matrix with size *r*. The phylogenetic covariance matrix (**C**) is a symmetric matrix of dimension *n* with diagonal elements equal to the sum of branch lengths from the root to each of the *n* tips and off-diagonals represent the sum of branch lengths from the root of the tree to the most recent common ancestor between each pair of tips (Felsenstein, 1973). Here, each of the **C**_*k*_matrices has only the sum of branch lengths which were assigned to the respective evolutionary rate matrix **R**_*k*_ (i.e., regime *k* painted on the tree). Thus, 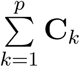 is equal to the phylogenetic covariance matrix (**C**) for the whole tree. Finally, *p* is the number of **R** matrix regimes fitted to the tree. When *p* is equal to 1, equation (1) reduces to the likelihood function for a single **R** matrix (Revell and Harmon, 2008).

### MCMC prior densities and sampling strategy

We have developed and implemented a Bayesian method to estimate one or more evolutionary rate matrices from phylogenetic comparative data. We model the prior density for the vector of root values (**a**) as an (improper) uniform or normal distribution and we use a uniform sliding window proposal density to sample the root value for every trait simultaneously. In contrast, the prior density and sampling scheme for the evolutionary rate matrix requires more elaboration because variance-covariance matrices are positive definite and are relatively hard to be estimated. We model **R** with two independent distributions; one for the standard deviations and another for the correlation matrix (Barnard et al., 2000; Zhang et al., 2006). At each step of the MCMC the proposals for the standard deviations of each trait and the correlation matrix are carried independently and **R** is reconstructed from these elements in order to evaluate the likelihood function of the model (Caetano and Harmon, 2017). This method allows the prior density for the rates (standard deviations) to be parametrized independently of the evolutionary integration (correlation matrix). Under this parametrization, one can assign any prior distribution of positive real values to the standard deviations (here we use an improper uniform or a exponential density as priors) and the prior for the correlation matrix is modelled as the Cholesky decomposition of variance-covariance matrices sampled from an inverse-Wishart distribution (Zhang et al., 2006; Caetano and Harmon, 2017). This parameter extension approach is named ‘separation-strategy’ (Barnard et al., 2000; Zhang et al., 2006) because it relies on the independent modelling of the vector of standard deviations and the correlation matrix that together compose the evolutionary rate matrix. The advantage of the separation-strategy is twofold; it allows for intuitive modelling of rates of evolution and evolutionary integration and it is an efficient proposal scheme, because matrices are guaranteed to be positive definite at every draw (Barnard et al., 2000; Zhang et al., 2006).

### A new pruning algorithm for multivariate Brownian motion with multiple regimes

The likelihood function for the evolutionary rate matrix as shown in equation 1 requires the matrix inversion and determinant to be computed for the sum of the Kronecker product between each **R_*k*_** and **C_*k*_**matrices. However, the matrices resulted from this product can be very large because each **R** has dimension equal to the number of traits in the data whereas **C** is as large as the number of tips in the phylogeny. Some methods can be used to speed up the computation in the case of multiple rate regimes applied to the tree. For instance, the ‘rpf’ method avoids the explicit computation of the matrix inversion and determinant by applying Cholesky factors (Gustavson et al., 2010; Clavel et al., 2015) whereas Goolsby (2016) recently introduced the use of pairwise composite likelihoods, which consists of the product of the pairwise likelihoods computed for all combinations of traits (but see Adams and Collyer, 2017). These methods reduce the computational time for the evaluation of the likelihood but are still more time consuming than the pruning algorithm (Felsenstein, 1973; Freckleton, 2012; Caetano and Harmon, 2017). Here, we expand the pruning algorithm applied to the multivariate Brownian motion model (Felsenstein, 1973; Freckleton, 2012) to compute the likelihood even when multiple evolutionary rate matrices are fitted to different branches of the phylogenetic tree. A detailed explanation of the algorithm is in the Supplementary Material. The algorithm is implemented in the R package ratematrix (Caetano and Harmon, 2017) and we also provide an example with commented code in the online appendix (http://dx.doi.org/XXXX/dryad).

### Incorporating uncertainty in regime configurations and phylogenetic trees

Our approach can integrate multiple evolutionary rate matrix regimes fitted to the same phylogenetic tree. Rate regimes are often informed by a predictive categorical trait which states are hypothesized to be associated with shifts in the tempo and mode of evolution of the traits under study. Comparative methods often map such regimes to phylogenetic trees applying an ancestral reconstruction approach. For intance, Huelsenbeck et al. (2003) described the stochastic character mapping method which utilizes a transition rate matrix (**Q**) for the states of a discrete trait to simulate character evolution histories across the branches of the phylogenetic tree. A collection of simulated histories using stochastic mapping based on an estimated **Q** matrix represents the uncertainty associated with the reconstruction of the discrete states on the phylogeny, both in the number of changes as well as the position of the regimes on the branches of the tree (see Fig. 1 steps 1-3). Unfortunatelly, most phylogenetic comparative methods do not directly incorporate such uncertainty in their estimates and users are required to repeat the analysis for each regime configuration on a sample of stochastic maps.

In order to facilitate incorporation of uncertainty in ancestral state estimates we implemented a MCMC that integrates over multiple rate regime configurations (Fig. 1). At each step of the MCMC one regime configuration is randomly sampled from a pool of stochastic maps and is used to evaluate the likelihood of the model. Thus, the MCMC chain visits all the stochastic mapped histories in the pool and incorporates the information about multiple ancestral reconstructions from this sample of character histories in the posterior distribution of parameter estimates for the model (Fig. 1 step 4). This approach assumes that each regime configuration in the pool has equal chance to be sampled. However, one can assign a sampling frequency to each of the regime configurations on the pool as a result of some previous analysis, such as the density of a posterior distribution of stochastic maps. This same approach can be used to incorporate uncertainty in phylogenetic estimation if the pool is comprised of samples of phylogenetic trees from a posterior distribution or bootstrap analyses. Although this method is able to incorporate the uncertainty related to alternative regime configurations, different topologies and sets of branch lengths, it is not a joint estimation of the tree and the model because the MCMC applies proposal steps exclusively to the parameters of the trait evolution model.

### Testing for shifts between rate regimes

A useful criterion to perform model selection in a Bayesian framework is the Bayes factor (Kass and Raftery, 1995), which is a ratio between the marginal likelihoods of the competing models. However, the estimation of marginal likelihoods is a computationally expensive and contentious task. One of the most accurate methods to estimate the marginal likelihood is the stepping stone approach. This method consists of taking samples from a series of weighted posterior distributions by scaling the likelihood of the model so that a continuum between the prior and the posterior is created (Fan et al., 2011; Xie et al., 2011; Uyeda and Harmon, 2014). However, the stepping stone method adds significantly to the computation burden of the analysis, because each step of the continuum represents a complete MCMC chain and a large number of steps are required to produce a sufficient approximation of the marginal likelihood (Uyeda and Harmon, 2014).

Here, we do not use Bayes factor to compare models, although implementation is feasible for future work. We focus our interpretation of results on the distribution of posterior parameter estimates, and quantify the amount of uncertainty and the magnitude of the difference between components of the evolutionary rate matrices fitted to different regimes of the tree. We implemented summary statistics that provide a framework to decide whether there is enough signal in the data to support a model comprised by multiple **R** matrix regimes. We check for shifts in the rates of evolution of each individual trait and in the structure of evolutionary correlation among traits. Then, we perform tests by computing the difference between summary statistics sampled from the joint posterior distribution of rate regimes and checking for an overlap smaller than 2.5% between distributions which indicates rate regimes are likely samples from distinct distributions. We apply these tests for each element of the evolutionary rate matrix, the correlation matrix as well as the rates of evolution for each trait, resulting in a full picture of the macroevolutionary pattern across the phylogenetic tree. Studying integration between each trait can be more informative than using an average measure of evolutionary correlation across the whole data, because the way integration is distributed among traits can have an important impact on trait disparity through time (Gerber, 2013).

### Simulation study

We performed extensive simulations to test the performance of our Bayesian MCMC estimates and the use of summary statistics under different scenarios of correlated and uncorrelated evolution varying both the size of the phylogeny and the number of traits. We used rejection sampling to simulate phylogenetic trees under a homogeneous birth-death model containing at least one monophyletic clade comprised by a fixed number of lineages for each simulation scenario. Then we generated data sets with two multivariate rate regimes and set the shift at the base of the focus clade (Fig. S1). We used data sets with 3 and 6 traits and a large variation of tree sizes, from small trees with 25 species to fairly large trees of 1000 tips. Tables 1 and 2 show the combinations of tree size, focus clade size, and number of traits used in our simulation study. For each combination of tree size and data set size we tested 8 distinct scenarios of multivariate evolution: no shift between regimes (Single), change from positive to negative evolutionary correlations (Orient), weak shift of rates (Rates I), strong shift of rates (Rates II), weak shift in the strength of correlation among all traits (Integ I), strong shift in the strength of correlation among all traits (Integ II), a weak shift on the correlation pattern on part of the traits (Pattern I), and a strong shift on the correlation pattern on part of the traits (Pattern II). Figures S1 and S2 show the parameter values used to simulate the data for each treatment using 3 and 6 traits, respectively.

**Table 1:** Proportion of simulation replicates showing support for two **R** matrix regimes under likelihood ratio test (LRT) and using summary statistics computed from the posterior distribution. We accepted a difference between rate regimes if at least one of the elements of the matrix showed overlap between the posterior estimates for the regimes lower than 2.5%. The ‘rates’ summary statistics compares the rates of evolution whereas ‘correlation’ contrasts the structure of evolutionary correlation between regimes. Accuracy was measured as the proportion of parameter estimates for which the true value were within the 95% highest posterior density. Simulations were performed under eight scenarios of correlated evolution as showed in Figure S1: no shift (Single), shift of orientation (Orient), weak (Rates I) and strong (Rates II) shift of rates, weak (Integ I) and strong (Integ II) shift of integration, and weak (Pattern I) and strong (Pattern II) shift of pattern.

|  | Single | Orient | Rates I | Rates II | Integ I | Integ II | Pattern I | Pattern II |
| --- | --- | --- | --- | --- | --- | --- | --- | --- |
| 25 species (6 in focal clade) and 3 traits |  |  |  |  |  |  |  |  |
| LRT | 0.16 | 0.57 | 0.37 | 0.82 | 0.17 | 0.35 | 0.21 | 0.21 |
| rates | 0.09 | 0.04 | 0.38 | 0.78 | 0.06 | 0.05 | 0.04 | 0.06 |
| correlation | 0.01 | 0.36 | 0.00 | 0.04 | 0.01 | 0.07 | 0.01 | 0.03 |
| accuracy rates | 0.94 | 0.94 | 0.94 | 0.95 | 0.94 | 0.94 | 0.94 | 0.95 |
| accuracy correlation | 0.97 | 0.97 | 0.98 | 0.96 | 0.98 | 0.98 | 0.98 | 0.98 |
| 50 species (12 in focal clade) and 3 traits |  |  |  |  |  |  |  |  |
| LRT | 0.13 | 0.81 | 0.53 | 0.97 | 0.11 | 0.50 | 0.18 | 0.13 |
| rates | 0.06 | 0.06 | 0.61 | 0.98 | 0.04 | 0.09 | 0.08 | 0.05 |
| correlation | 0.06 | 0.86 | 0.04 | 0.10 | 0.06 | 0.36 | 0.07 | 0.10 |
| accuracy rates | 0.95 | 0.95 | 0.94 | 0.96 | 0.93 | 0.93 | 0.94 | 0.95 |
| accuracy correlation | 0.96 | 0.96 | 0.96 | 0.96 | 0.97 | 0.96 | 0.96 | 0.98 |
| 100 species (25 in focal clade) and 3 traits |  |  |  |  |  |  |  |  |
| LRT | 0.04 | 0.99 | 0.79 | 1.00 | 0.18 | 0.79 | 0.28 | 0.37 |
| rates | 0.01 | 0.04 | 0.81 | 1.00 | 0.07 | 0.07 | 0.12 | 0.05 |
| correlation | 0.03 | 1.00 | 0.04 | 0.06 | 0.13 | 0.75 | 0.12 | 0.39 |
| accuracy rates | 0.96 | 0.96 | 0.95 | 0.96 | 0.95 | 0.95 | 0.93 | 0.95 |
| accuracy correlation | 0.98 | 0.96 | 0.95 | 0.95 | 0.96 | 0.95 | 0.96 | 0.95 |
| 200 species (50 in focal clade) and 3 traits |  |  |  |  |  |  |  |  |
| LRT | 0.11 | 1.00 | 0.98 | 1.00 | 0.25 | 0.99 | 0.21 | 0.76 |
| rates | 0.09 | 0.06 | 1.00 | 1.00 | 0.07 | 0.08 | 0.06 | 0.04 |
| correlation | 0.04 | 1.00 | 0.03 | 0.03 | 0.25 | 1.00 | 0.27 | 0.72 |
| accuracy rates | 0.97 | 0.95 | 0.94 | 0.94 | 0.95 | 0.93 | 0.94 | 0.95 |
| accuracy correlation | 0.96 | 0.96 | 0.95 | 0.95 | 0.96 | 0.96 | 0.96 | 0.93 |

**Table 2:** Proportion of simulation replicates showing support for two **R** matrix regimes under likelihood ratio test (LRT) and using summary statistics computed from the posterior distribution. Maximum likelihood estimates for phylogenetic trees larger than 200 tips were not performed due to excessive computational burden. We accepted a difference between rate regimes if at least one of the elements of the matrix showed overlap between the posterior estimates for the regimes lower than 2.5%. The ‘rates’ summary statistics compares the rates of evolution whereas ‘correlation’ contrasts the structure of evolutionary correlation between regimes. Accuracy was measured as the proportion of parameter estimates for which the true value were within the 95% highest posterior density. Simulations were performed under eight scenarios of correlated evolution as showed in Figure S2: no shift (Single), shift of orientation (Orient), weak (Rates I) and strong (Rates II) shift of rates, weak (Integ I) and strong (Integ II) shift of integration, and weak (Pattern I) and strong (Pattern II) shift of pattern.

|  | Single | Orient | Rates I | Rates II | Integ I | Integ II | Pattern I | Pattern II |
| --- | --- | --- | --- | --- | --- | --- | --- | --- |
| 200 species (50 in focal clade) and 6 traits |  |  |  |  |  |  |  |  |
| LRT | 0.07 | 1.00 | 1.00 | 1.00 | 0.57 | 1.00 | 0.63 | 1.00 |
| rates | 0.15 | 0.15 | 1.00 | 1.00 | 0.14 | 0.20 | 0.17 | 0.16 |
| correlation | 0.23 | 1.00 | 0.22 | 0.19 | 0.81 | 1.00 | 0.46 | 0.92 |
| accuracy rates | 0.91 | 0.93 | 0.90 | 0.91 | 0.93 | 0.90 | 0.91 | 0.92 |
| accuracy correlation | 0.95 | 0.96 | 0.94 | 0.95 | 0.95 | 0.95 | 0.95 | 0.94 |
| 400 species (100 in focal clade) and 6 traits |  |  |  |  |  |  |  |  |
| LRT | — | — | — | — | — | — | — | — |
| rates | 0.21 | 0.21 | 1.00 | 1.00 | 0.15 | 0.14 | 0.14 | 0.16 |
| correlation | 0.28 | 1.00 | 0.22 | 0.22 | 0.99 | 1.00 | 0.81 | 1.00 |
| accuracy rates | 0.93 | 0.92 | 0.94 | 0.94 | 0.94 | 0.93 | 0.92 | 0.94 |
| accuracy correlation | 0.94 | 0.94 | 0.96 | 0.96 | 0.95 | 0.95 | 0.96 | 0.94 |
| 500 species (200 in focal clade) and 6 traits |  |  |  |  |  |  |  |  |
| LRT | — | — | — | — | — | — | — | — |
| rates | 0.08 | 0.15 | 1.00 | 1.00 | 0.20 | 0.15 | 0.12 | 0.09 |
| correlation | 0.29 | 1.00 | 0.29 | 0.21 | 1.00 | 1.00 | 1.00 | 1.00 |
| accuracy rates | 0.94 | 0.94 | 0.94 | 0.94 | 0.93 | 0.94 | 0.94 | 0.94 |
| accuracy correlation | 0.95 | 0.95 | 0.94 | 0.95 | 0.96 | 0.95 | 0.95 | 0.95 |
| 1000 species (400 in focal clade) and 6 traits |  |  |  |  |  |  |  |  |
| LRT | — | — | — | — | — | — | — | — |
| rates | 0.00 | 0.00 | 1.00 | 1.00 | 0.00 | 0.00 | 0.17 | 0.00 |
| correlation | 0.00 | 0.60 | 0.00 | 0.00 | 1.00 | 1.00 | 0.40 | 0.40 |
| accuracy rates | 0.95 | 0.96 | 0.95 | 0.95 | 0.94 | 0.94 | 0.95 | 0.95 |
| accuracy correlation | 0.95 | 0.96 | 0.95 | 0.96 | 0.95 | 0.95 | 0.95 | 0.96 |

We estimated the posterior distribution for the evolutionary rate matrices of each simulation replicate using two independent MCMC chains of 1.5 million generations each and with starting points randomly sampled from the prior distribution. We checked for convergence using the potential scale reduction factor (Gelman and Rubin, 1992) as implemented in the package ratematrix (Caetano and Harmon, 2017). Then we used our set of summary statistics to ask if there is enough information in the data to support a shift in either the rates or the structure of correlated evolution among traits. Additionally, we computed the accuracy of estimates by checking the proportion of posterior distributions for each model parameter that contained the true value within the 95% highest posterior density (HPD). In order to check for congruence between our approach and maximum likelihood estimators, we used the R package mvMORPH (Clavel et al., 2015) to find the best model using likelihood ratio tests (one regime versus two regimes). The comparison between the likelihood ratio test and our posterior check approach is not a formal evaluation of model test performance, since the two approaches are fundamentally distinct. On the other hand, this serve as a pragmatic comparison to show whether we can adopt the use of summary statistics calculated from the posterior distribution to make reliable choices between models with direct incorporation of uncertainty in parameter estimates while retaining the explanation power of a formal model test approach. We did not performed maximum likelihood estimates with 6 traits and trees larger than 200 species due to the extreme computational burden without the usage of the pruning algorithm introduced herein (see Table 2).

One concern with multivariate models of trait evolution is about their performance when the number of traits becomes large with respect to the size of the phylogenetic tree (Adams and Collyer, 2017). If there are more traits or trait dimensions (*sensu* Adams, 2014b) than the number of tips in the phylogeny, the likelihood for the multivariate Brownian motion model cannot be computed (Adams, 2014b; Adams and Collyer, 2017; Caetano and Harmon, 2017). Similarly, as the number of traits or trait dimensions in the data increases relative to the number of species the power of the inference is expected to decrease (Adams and Collyer, 2017). Here we show the relationship between the degree of uncertainty in the parameter estimates (i.e., the width of the posterior distribution) and the accuracy (i.e., frequency of true values within the 95% HPD) in function of sample size. We define sample size as the number of tips in the phylogeny compared among datasets with the same number of traits (i.e., equal number of free parameters in the model). For this test we pooled the results from all simulations, but we separated estimates for the focus and background regimes as independent samples. This procedure is unlikely to bias our results, because the MCMC estimation for each regime was performed with independent proposal steps.

For all simulations we used an (improper) uniform prior for the vector of standard deviations, a marginally uniform prior for the correlation matrix (Barnard et al., 2000), and a multivariate normal prior for the vector of phylogenetic means centered in the mean of the tip data for each trait and with standard deviation equal to two times the standard deviation of the tip data (Fig. S3). We chose an informative prior for the phylogenetic mean in order to facilitate the convergence of the MCMC chains, since the root values are not the primary focus of this set of simulations. Nevertheless, we repeated a subset of the simulations using an uninformative prior assigned to the root values to show that the MCMC also performs well under this scenario. We checked for convergence using the Gelman and Rubin (1992) test applied to each parameter of the model (i.e., root values and each element of the evolutionary rate matrices). We simulated phylogenies, traits and mapped regimes using the R package phytools (Revell, 2012) and performed all parameter estimates with the package ratematrix (Caetano and Harmon, 2017). Simulated datasets and code to replicate results are available from the Dryad Digital Repository: http://dx.doi.org/XXXX/dryad.

### Empirical examples

We use two examples to show the performance of the approach with empirical datasets and to further explore the impact of the direct incorporation of uncertainty in parameter estimates and model comparison. The first example tests for a shift in the evolutionary integration among anoles traits during the Caribbean radiation. Then, we repeat the analysis from Revell and Collar (2009) study on the evolution of buccal traits in Centrarchidae fishes. Data, results, and code to replicate the analyses for both empirical examples are available from the Dryad Digital Repository: http://dx.doi.org/XXXX/dryad.

Anoles are small lizards that live primarily in the tropics. There are nearly 400 anole species with diverse morphology and they have become a model system for studies of adaptive radiation and convergence (Losos, 2009; Mahler et al., 2013; Poe et al., 2017, 2018, and references therein). The ancestral distribution of the genus is in Central and South America and the history of the clade includes island dispersion and radiation as well as dispersal back to the mainland (Nicholson et al., 2005; Losos, 2009; Poe et al., 2017). The adaptive radiation of anoles to the Caribbean islands and the repeated evolution of ecomorphs are the main focus of evolutionary studies in the genus (Mahler et al., 2010; Losos, 2009; Mahler et al., 2013). However, mainland anoles are distributed from the north of South America to the south of North America and show more species (60% of all species) than island anoles as well as equally impressive morphological diversity (Losos, 2009; Poe et al., 2017, 2018). Mainland and island anole species form distinct morphological clusters (Pinto et al., 2008; Schaad and Poe, 2010; Moreno-Arias and Calder´on-Espinosa, 2016), but rates of trait evolution have been shown not to be consistently different (Pinto et al., 2008; Poe et al., 2018). Island ecomorphs can be readily distinguished by body size and the morphology of limbs, head and tail (Losos, 2009; Mahler et al., 2013). Thus, it is plausible that a shift in the structure of evolutionary integration among those traits associated with the radiation to the islands played an important role on the exploration of novel regions of the morphospace and allowed the repeated evolution of specialized morphologies. Herein we test this hypothesis by fitting three evolutionary rate matrix regimes defined by lineages occupying South America mainland (ancestral distribution), Caribbean island species, and the descendents of the more recent dispersal back to the mainland. By comparing the patterns of trait evolution between the ancestral mainland lineages with the lineages that dispersed back to the mainland we can test whether the tempo and mode of evolution is convergent and support the hypothesis that changes in trait evolution are correlated with shifts between mainland and island distribution.

We compiled data for snout-vent length (SVL), tail length (TL), and head length (HL) of 147 anole species (98 Caribbean and 49 mainland species) made available by Mahler et al. (2013) and Moreno-Arias and Calder´on-Espinosa (2016). We chose this set of traits because they are important for niche partitioning among anoles (Pinto et al., 2008; Losos, 2009; Mahler et al., 2013) and also provided the best species coverage given the data currently available. We used Poe et al. (2017) dated maximum clade credibility tree (hereafter time tree) for all comparative analyses, but we trimmed the phylogeny to include only the species that we have morphological data. To map the different **R** matrix regimes to the time tree we classified species as ‘island’, ‘ancestral mainland’, or ‘dispersal mainland’. To differentiate between ancestral and dispersal mainland distributions we used results from the biogeographical reconstructions of Poe et al. (2017). We estimated the best model for transition rates between states assuming the most common ancestor for the genus was a mainland lineage (following Poe et al., 2017). Then we used the package ‘phytools’ (Revell, 2012) to perform 100 stochastic mapping simulations conditioned on the estimated transition matrix. We set the model to estimate one **R** matrix for each mapped state (‘island’, ‘ancestral mainland’, or ‘dispersal mainland’) and we used the pool of 100 stochastic maps in the MCMC to take into account the uncertainty associated with ancestral state estimation. We ran four independent MCMC chains of 2 million generations each and used a random sample from the prior as the starting point of each chain. We set an (improper) uniform prior for the phylogenetic mean and the vector of standard deviations and a marginally uniform prior on the correlation matrices. We discarded 25% of each MCMC chain as burn-in and checked for convergence using the potential scale reduction factor (Gelman and Rubin, 1992).

In addition to the analyses of mainland and island anoles, we replicated the study by Revell and Collar (2009) as an exercise to contrast the inference of evolutionary rate matrices in the presence of a direct estimate of uncertainty provided by the posterior densities. Revell and Collar (2009) showed that the evolution of a specialized piscivorous diet in fishes of the genus *Micropterus* is associated with a shift towards a stronger evolutionary correlation between buccal length and gape width (see Fig. 1 in Revell and Collar, 2009). This tighter integration might have allowed *Micropterus* lineages to evolve better suction feeding performance. For this analysis we used the same data and phylogenetic tree made available by the authors. We set prior distributions using the same approach for the analysis of anole lizards described above. We also ran four MCMC chains starting from random draws from the prior for 1 million generations and checked for convergence using the potential scale reduction factor (Gelman and Rubin, 1992). Finally, we compared the result of the posterior distribution of parameter estimates with the MLE by estimating a confidence interval for the MLE using parametric bootstraps.

## Results

### Performance of the method

We estimated the posterior distribution of evolutionary rate matrices for 6,400 simulated data sets distributed among multiple scenarios of correlated evolution while exploring the effect of number of traits and size of the phylogenetic tree on the performance of the model. Tables 1 and 2 show detailed results for all simulation replicates whereas figures 2 and 3 show examples of the posterior distribution for each simulation scenario together with vertical lines pointing to the true values used to simulate the data. In overal, our results show that the use of the summary statistics computed from the posterior distribution of parameter estimates is a reliable method to study rates of evolution as well as the pattern of evolutionary correlation among traits.

**Figure 2:**
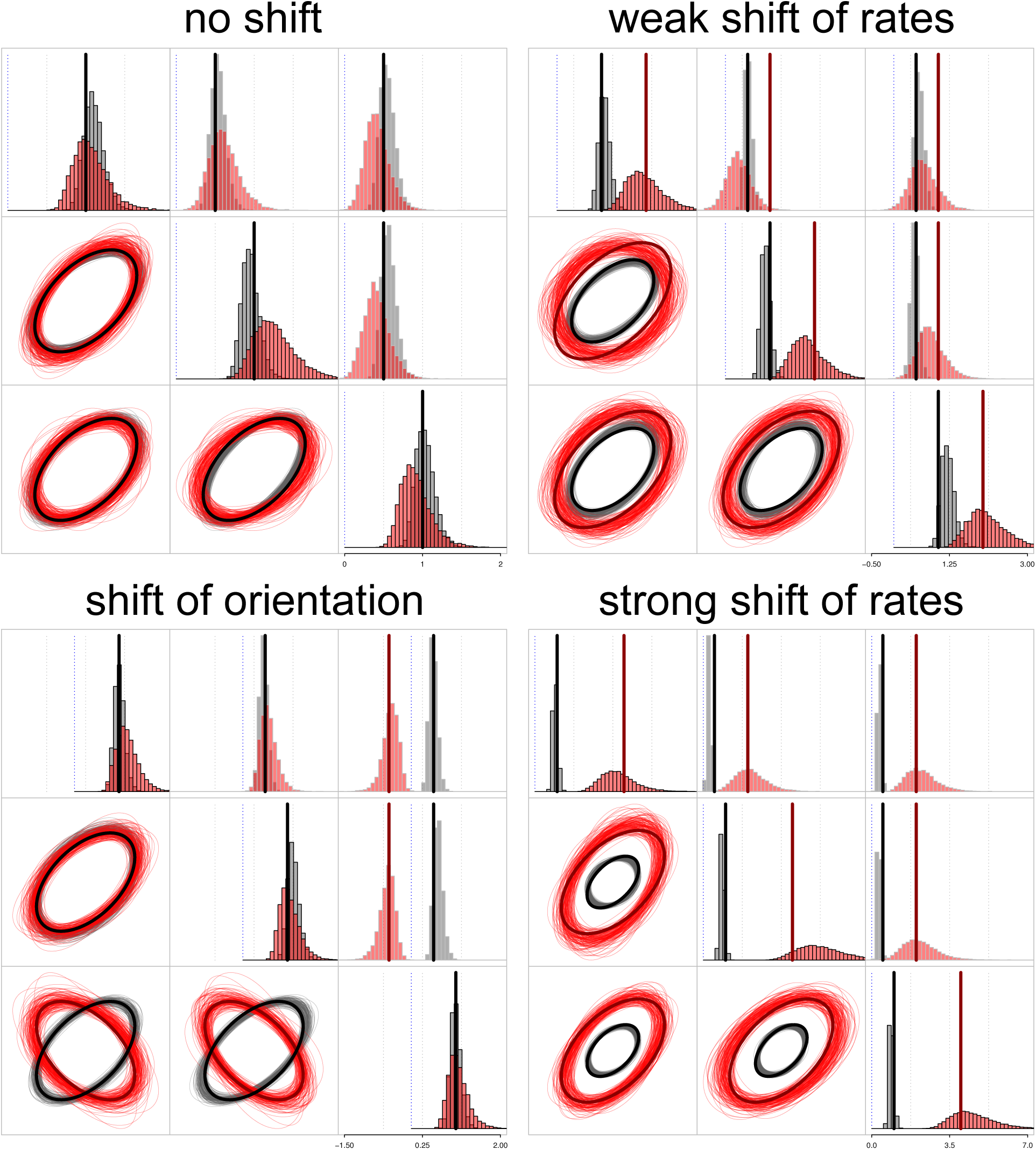
Example of posterior distribution for the first four simulation treatments with three traits and 200 species. Top-left plot shows results with no shift in the evolutionary rate matrix regime and bottom-left plot shows a shift in the orientation of the evolutionary correlation among traits. Top-right and bottom-right plots show weak and strong shifts in the rates of correlation, respectivelly. Estimates for the background regime are showed in gray and for the focus regime in orange. Broad vertical lines and ellipse lines show the true parameter values used for the simulations (see also Figure S1). For each plot: diagonal histograms show evolutionary rates for each trait, upper-diagonal histograms show pairwise evolutionary covariation, and lower-diagonal ellipses are samples from the posterior distribution showing the 95% confidence interval of each bivariate distribution. Table 1 shows the aggregate results for each simulation treatment.

**Figure 3:**
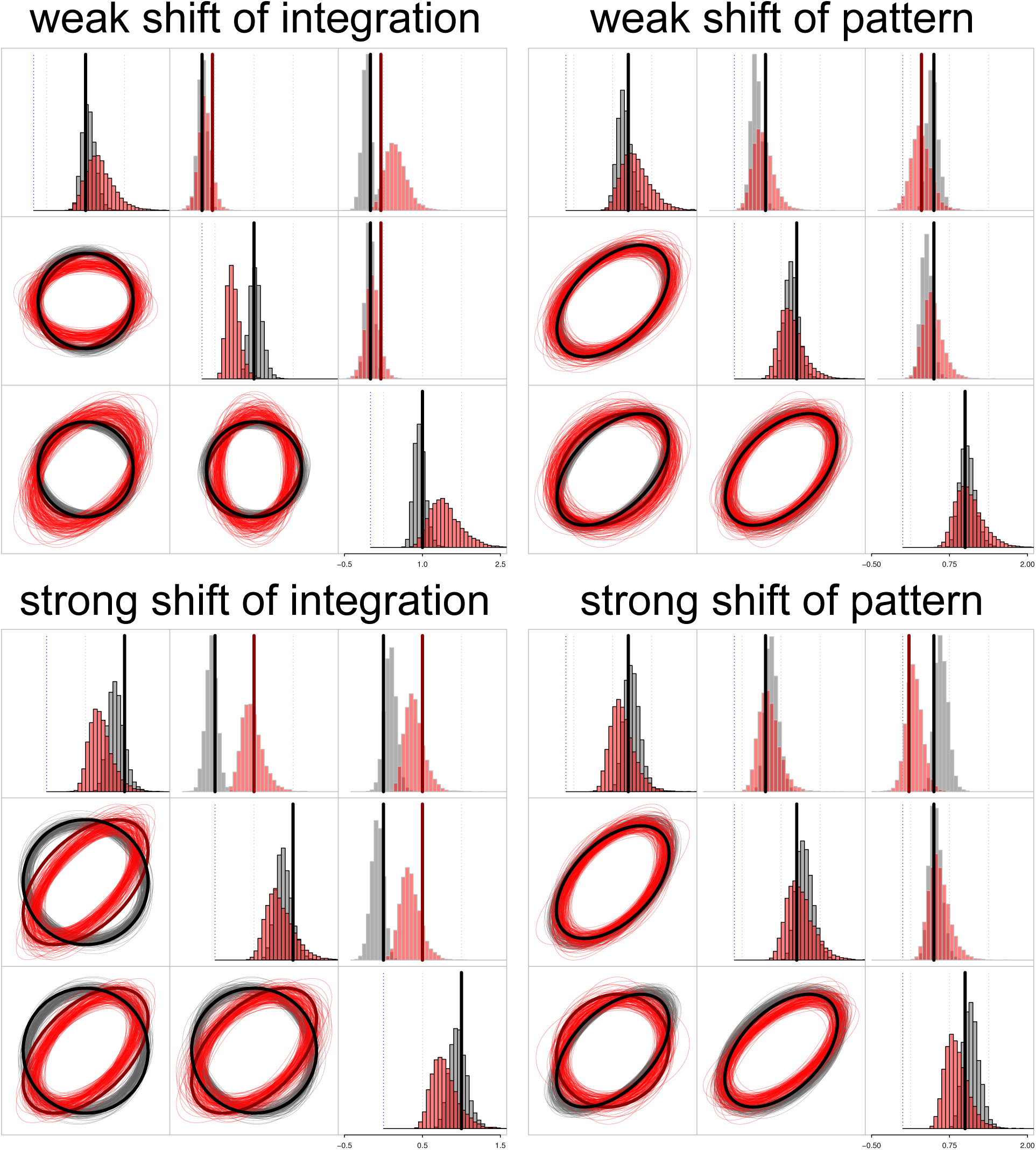
Example of posterior distribution for the last four simulation treatments with three traits and 200 species. Top-left and bottom-left plots show results with weak and strong shifts of integration among traits. Top-right and bottom-right show weak and strong shifts of the pattern of evolutionary correlation among the traits. Estimates for the background regime are showed in gray and for the focus regime in orange. Broad vertical lines and ellpse lines show the true parameter value used for simulations (see also Figure S1). For each plot: diagonal histograms show evolutionary rates for each trait, upper-diagonal histograms show pairwise evolutionary covariation, and lower-diagonal ellipses are samples from the posterior distribution showing the 95% confidence interval of each bivariate distribution. Table 1 shows the aggregate results for each simulation treatment.

When we simulated data using a single multivariate rate of evolution across all branches of the tree (treatment ‘Single’) and 3 traits, the results based on the summary statistics computed from the posterior distribution of parameter estimates showed between 1% and, at the worst case scenario, 9% of replicates with enough signal to wrongly distinguish between **R** matrices fitted to different regions of the tree (Table 1). In contrast, likelihood ratio tests (LRTs) applied to the same data showed error rates almost two times higher (Table 1). Results with 6 traits were only acceptable with phylogenetic trees larger than 500 species (Table 2). Smaller phylogenetic trees resulted in elevated error rates, suggesting that large sample sizes are required to jointly estimate the rates and the evolutionary correlation among many traits. The performance of LRTs were especially lower in cases of reduced relative number of species per trait in the data (see Tables 1 and 2).

Simulations with 3 traits and a shift in rates while holding the pattern of evolutionary correlation among traits constant (Table 1 columns ‘Rates I’ and ‘Rates II’) showed an increase in power as the relative sample size increased. Phylogenies with 25 and 50 species resulted in about 1.5 to 2 times more replicates that detected a shift in rates when the focus clade was simulated with a four-fold increase in rates (‘Rates II’) compared to a two-fold change (‘Rates I’). However, almost all simulation replicates showed a strong signal for a shift in the rates of evolution for trees with 100 and 200 tips, suggesting the effect of tree size reached a plateau (Table 1). In the case of data sets with 6 traits, the power to detect shifts in rates of evolution is high across all tree sizes (Table 2). The evolutionary correlation statistics correctly showed low proportion of support for a change in the evolutionary correlation with 3 traits (Table 1) whereas results with 6 traits only reflected the true parameters with trees of large size (Table 2).

When changes in the pattern of integration were simulated in the data, results with 3 traits followed the same pattern as the shifts in rates of evolution—power increased with relative sample size (Table 1). However, weak changes in integration (‘Integ I’) and pattern (‘Pattern I’) showed the lowest number of simulation replicates detecting a shift (Table 1). In the case of 3 traits, the number of replicates that successfully detected a shift in the mode of evolution was up to four times higher in the simulation scenarios with a strong shift (‘Integ II’ and ‘Pattern II’). Results with 6 traits showed a different pattern. The power to detect shifts of integration among traits was high under all treatments and tree sizes, but smaller trees wrongly detected shifts in the rate of evolution between regimes more often (Table 2). In overal, the performance of the model was best in the scenario of shifts in orientation (Tables 1 and 2 column ‘Orient’) in which there is a strong change in the pattern of evolution, with some of the traits shifting from a positive to a negative correlation with other traits.

The accuracy of the MCMC estimator, measured as the percent of replicates that contained the true parameter values within the 95% highest posterior density (HPD) interval, was about 95% for both rates and evolutionary correlation among traits across all simulation replicates. This result is expected if parameters are correctly estimated with a MCMC (Cook et al., 2006). The accuracy of parameter estimates do not reflect the frequency in which our summary statistics correctly detected a shift. We computed summary statistics that respond to the width of the posterior distribution. When the uncertainty in parameter estimates increases the width of the posterior increases as well. However, the true value for each parameter still lies within the 95% HPD interval. For instance, the plot of the width of the posterior distributions in function of sample size, using the same simulation results, shows that the accuracy of parameter estimates increases with larger phylogenetic trees (Figure S4).

We used a multivariate normal prior for the root values centered at the mean of the tip data for all simulation replicates. The posterior distributions are showed on Figure S5. Changing the prior distribution from multivariate normal to uniform showed no detectable bias in the posterior distribution of root values under all simulation scenarios (Fig. S6).

### Empirical examples

We estimated the evolutionary rate matrix (**R**) for anole lineages by fitting three distinct regimes to the phylogenetic tree. We made a distinction between the ancestral mainland distribution and the more recent dipersal back to the mainland by fitting two independent regimes (Figure 5). Our results show no difference in the rates of evolution of traits among regimes, but lineages that dispersed back to mainland have the most distinct estimates of rates of evolution relative to the Caribbean species (Table S1). Island species seem to show faster morphological differentiation when compared to the clade that dispersed back to mainland (see Figure 5), which corroborates results reported by Poe et al., 2018. However, the uncertainty associated with posterior parameter estimates casts doubt on such conclusion. Unfortunately, further comparisons to previous results are difficult due to the absence of reported confidence intervals for parameter estimates. Likewise, there is no difference in the pattern of evolutionary correlation between regimes for most of the traits. Congruent with our results for evolutionary rates, the most distinct estimates for evolutionary correlation were between recent mainland dispersers and island species. The summary statistics computed from the posterior distribution indicates a shift in the correlation between body size and head length of island species and mainland dispersers (Table S1). Although traits show positive evolutionary correlation with body size, head length in mainland dispersers shifted towards a less tight relationship with body size.

In the case of the Centrarchidae fishes, there is a clear distinction between the results of maximum likelihood and Bayesian estimate for the evolutionary rate matrix regimes (Fig. 6). Under MLE, we found a significant difference between the **R** matrix regimes using likelihood ratio tests. There is a stronger evolutionary correlation between the gape width and the buccal length of the *Micropterus* clade (r=0.83) when compared with other lineages (r=0.36). In contrast, the direct incorporation of uncertainty in parameter estimates reveal an important overlap between the posterior densities for the **R** matrices estimated for each regime (Fig. 6). The posterior density does not show evidence of a shift towards stronger evolutionary correlation between gape width and buccal length in *Micropterus* (regime overlap in evolutionary correlation = 0.46, Fig. S8). Rates of evolution for both traits are also not different between the two regimes (ss-rates for gape width: 0.19, and buccal length: 0.08). Results from parametric bootstrap simulations, which estimate the uncertainty associated with maximum likelihood estimates, are congruent with the posterior distribution for the same model and show a similar degree of uncertanty (Fig. S9). Thus, after taking the uncertainty in parameter estimates from the Bayesian and the maximum likelihood analyses into account, it is unlikely that a shift on the pattern of evolutionary correlation happened in the *Micropterus* clade.

## Discussion

Here we implemented a Bayesian Markov chain Monte Carlo estimate of the evolutionary rate matrix. Our approach allows multiple regimes to be fitted to the same phylogenetic tree and integrates over a sample of trees or regime configurations to account for uncertainty in phylogenetic inference and ancestral state estimates. We also implement summary statistics to compare the posterior distribution of parameter estimates for different regimes. We show that our approach has good performance over a series of different scenarios of evolutionary integration when the size of the phylogenetic tree is large enough given the number of traits in the data set. The use of maximum likelihood estimate is definitely faster than Bayesian inference, since the MCMC chain requires many more evaluations of the likelihood function. However, our new extension of Felsenstein (1973) pruning algorithm applied when multiple **R** matrices are fitted to the same tree reduces the computation time of the likelihood for the model significantly (see Fig. 4 in Caetano and Harmon, 2017). For instance, the analyses including 6 traits fitted to large phylogenetic trees that we performed here would be virtually impossible without using our prunning algorithm. The integration of uncertainty in parameter estimates provided by the posterior distribution and the use of summary statistics to describe patterns in the data that can be directly relevant to our biological predictions are significant rewards for the longer time invested in data analysis when compared with MLE approaches.

**Figure 4:**
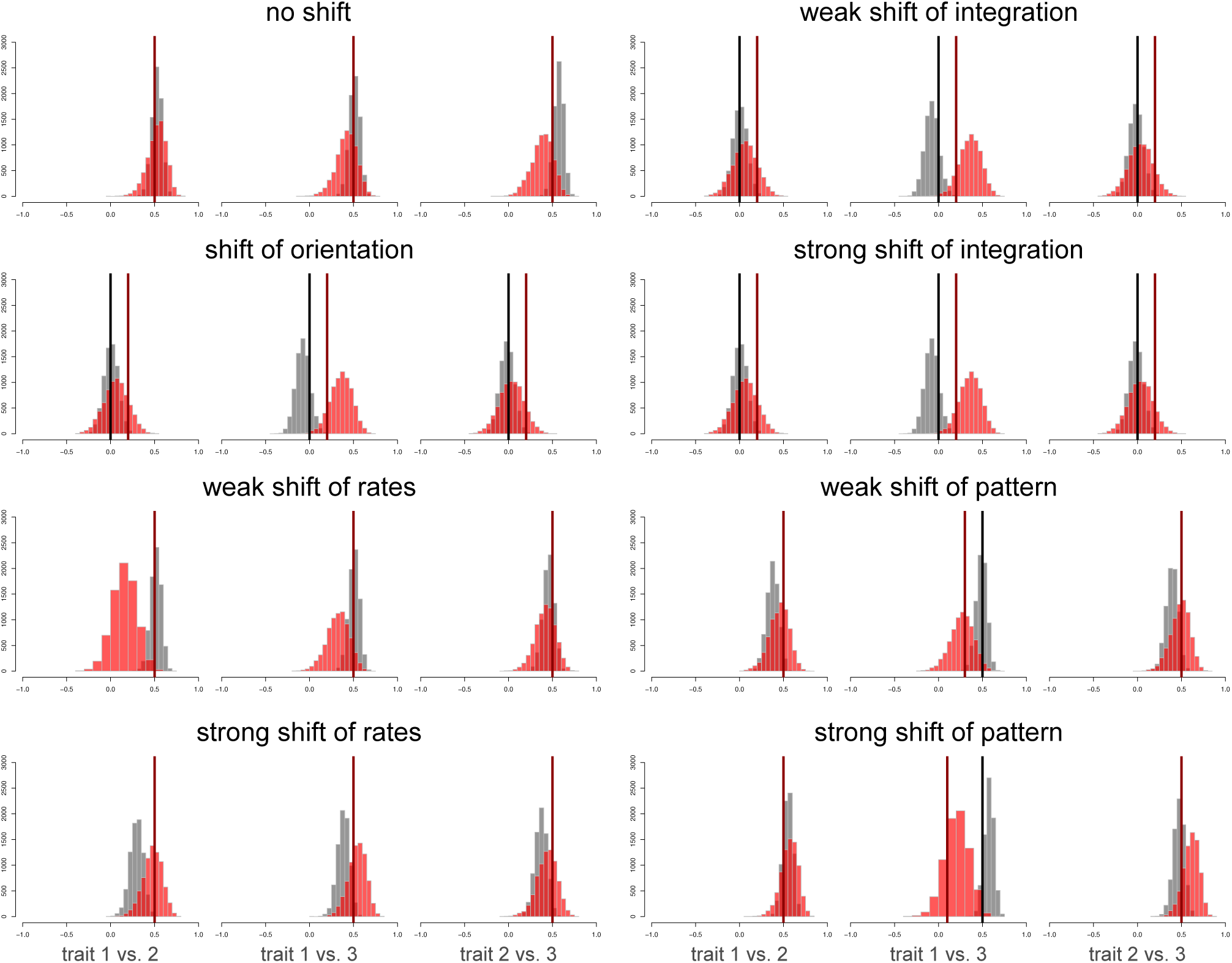
Example of posterior distribution for the pairwise evolutionary correlation among traits for the eight simulation treatments. Estimates for the background regime are showed in gray and for the focus regime in orange. Vertical lines show the true parameter values of the evolutionary correlations used for the simulations (see Figure S1). Table 1 shows the aggregate results for each simulation treatment.

**Figure 5:**
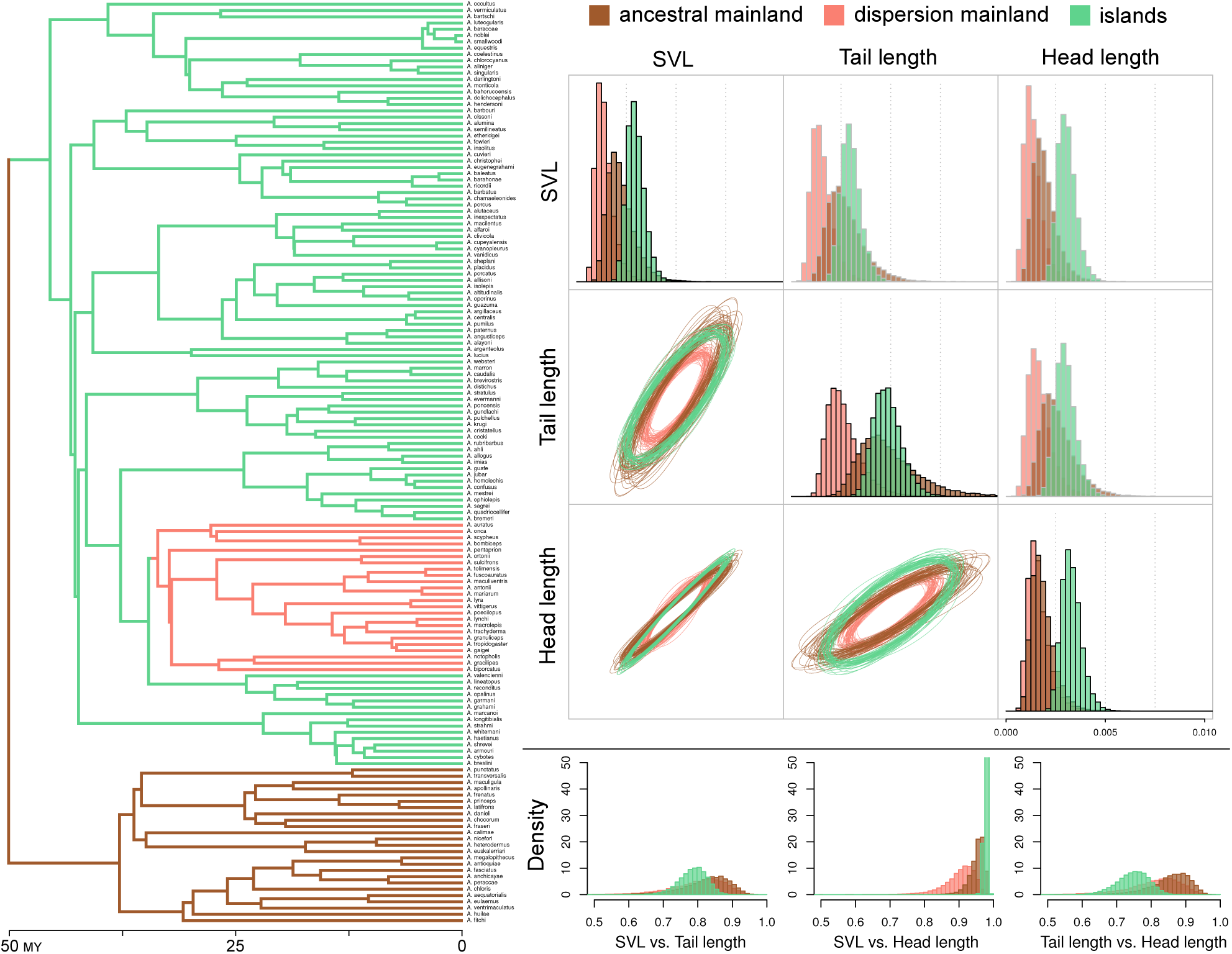
Posterior distribution of **R** matrix regimes fitted to the mainland anole, separated between ancestral mainland (brown) and the recent dispersal back to the mainland (salmon), and island anole (green) lineages. Left figure shows the maximum clade credibility tree (MCC) from Poe et al. (2017) with only the taxa used in this study. State reconstruction for the branches was performed with a stochastic map simulation using mainland as the root state for the genus and are congruent with original study (Poe et al., 2017). Right upper plot shows the posterior distribution of parameter estimates for the evolutionary rate matrices. Diagonal plots show evolutionary rates (variances) for each trait, upper-diagonal plots show pairwise evolutionary covariation (covariances), and lower-diagonal are samples from the posterior distribution of ellipses showing the 95% confidence interval of each bivariate distribution. Right bottom figure shows posterior density plots for the evolutionary correlation among traits.

**Figure 6:**
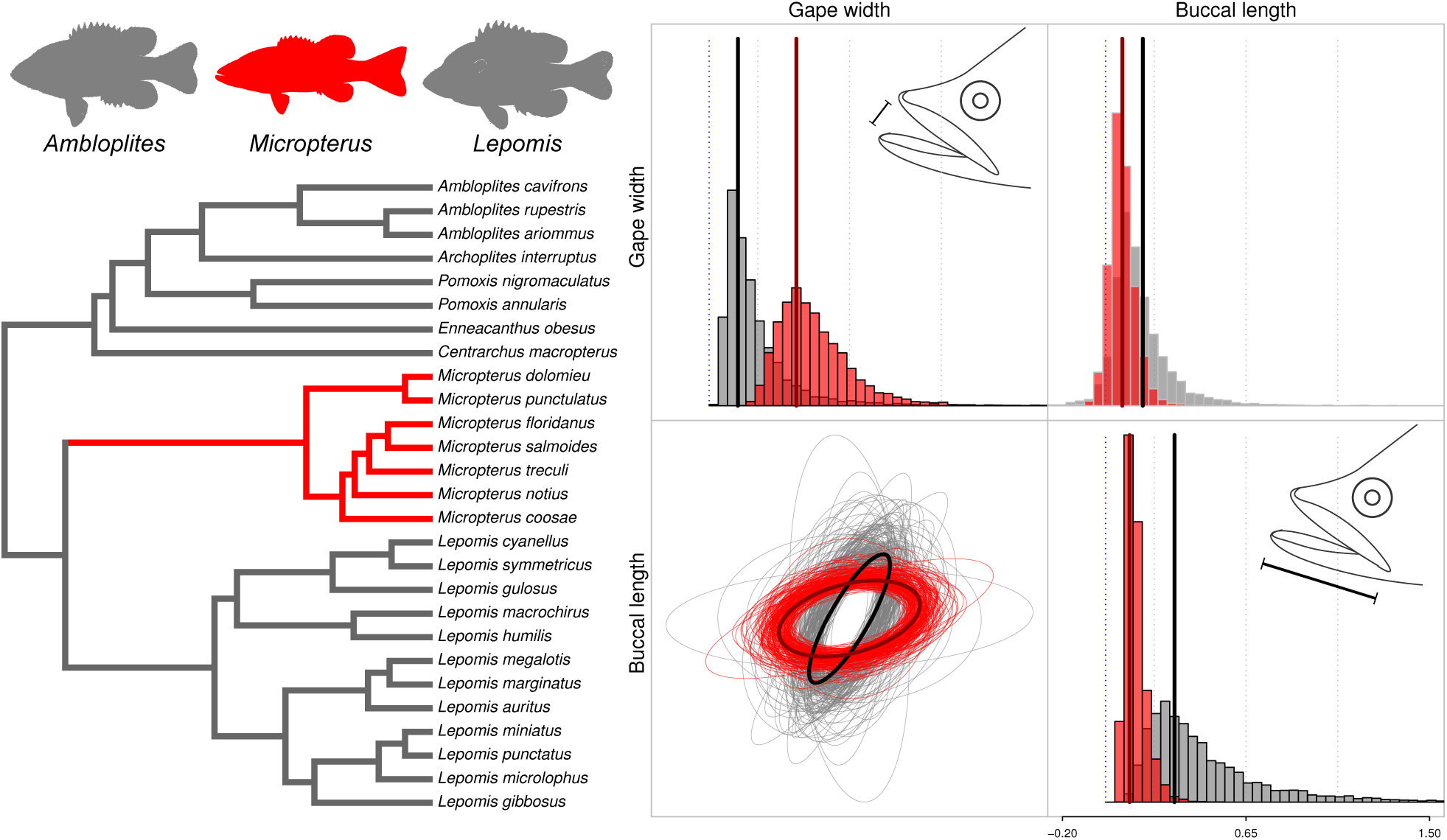
Posterior distribution of the **R** matrix regimes fitted to the background group (gray) and to the *Micropterus* clade (red). Left figure shows the phylogeny from (Revell and Collar, 2009) and the silhouette of some representatives of the Centrarchidae genera. Right plot shows the posterior distribution of parameter estimates for the evolutionary rate matrices. Diagonal plots show evolutionary rates (variances) for each trait, upper-diagonal plots show pairwise evolutionary covariation (covariances), and lower-diagonal are samples from the posterior distribution of ellipses showing the 95% confidence interval of each bivariate distribution. Broad vertical lines and ellipse lines matching the regime colors show the point estimate for the evolutionary rate matrices using maximum likelihood. See Figure S8 for the posterior distribution of evolutionary correlation between the traits.

The use of summary statistics to evaluate the overlap between the posterior distributions of parameter estimates from different regimes is an intuitive and reliable framework to make decisions of whether or not the data show a strong signal for multiple regimes. Our simulations showed that results from this approach are, in average, congruent with the likelihood ratio test. More importantly, summary statistics computed from the posterior distribution can recognize meaningful discrepancies between distinct evolutionary rate matrix regimes across a series of simulation scenarios. In this study we focused on the evolutionary rates for each trait and the evolutionary correlation among traits, but any other summary statistics computed over the posterior distribution of parameter estimates and representing an attribute of the model relevant for a given question could be implemented. For example, characteristics of the eigen-structure of the matrices or more formal tests such as the Flury hierarchy (Phillips and Arnold, 1999) or likelihood approaches to study modularity (Goswami and Finarelli, 2016) could be also implemented. This framework is flexible, does not require an estimate of the marginal likelihood and can be easily tailored towards specific biological predictions of the study system. On the other hand, it is important to note that the use of summary statistics does not constitute a formal model test, but instead asks the question of whether the parameter estimates for the regimes are distinct enough for us to accept the hypothesis of a change in the tempo and mode of trait evolution.

Recently, Adams and Collyer (2017) reviewed comparative approaches to study the evolution of multivariated data. They evaluated the performance and correcteness of methods using likelihood (Revell and Harmon, 2008; Revell and Collar, 2009; Clavel et al., 2015; Caetano and Harmon, 2017), pseudo-likelihood (Goolsby, 2016), and the likelihood-free algebraic generalization (Adams, 2014b). They show the evolutionary rate matrix is a reliable model when the ratio of the number of species in the phylogeny over the number of traits is suficiently large, which is congruent with our results. In fact, this is a common result for any statistical model using likelihood and is not specific to multivariate models, since parameter estimates are more accurate as the sample size increases. Phylogenetic comparative models are powerful tools, but there is a limit to how much we can learn from trees of small size and this limitation have been widely recognized (e.g., Beaulieu et al., 2012; Davis et al., 2013; Uyeda and Harmon, 2014; Rabosky and Goldberg, 2017). Moreover, our pruning algorithm solves the issue of ill-conditioned covariance matrices pointed out by Adams and Collyer (2017) and allows for data sets larger than previous implementations.

Adams and Collyer (2017) failed to distinguish between data sets comprised by multiple traits and single traits composed of multiple dimensions, although this distinction have been made clear in previous studies (Adams, 2014b,a; Adams and Collyer, 2015). This is an important concept because 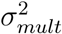 (Adams, 2014b) computes a single net evolutionary rate for all traits and ignores (by factoring out of the estimate) the evolutionary correlation among traits. Thus, 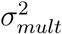 is not recommended if different traits in the data can show distinct rates of evolution or if the structure of evolutionary correlation among traits is of interest. Likelihood-based models, such as the one described here, are the correct choice to estimate evolutionary patterns among multiple traits. The algebraic generalization (Adams, 2014b) should be restricted to highly-dimensional traits such as shape data.

Point estimates such as the maximum likelihood can generate a false impression of certainty that may limit our biological interpretations if not accompanied by estimates of the variance. It is possible to calculate the confidence interval around the MLE and use this interval to check for overlaps in the parameter estimates (i.e., using the Hessian matrix). The disadvantage of this approach is that the confidence interval provides only the percentiles of the density around the MLE. Another alternative is the implementation of parametric bootstraps. However, these often take more time to estimate than the posterior distribution and are much less flexible approaches. Maximum likelihood estimates of models of trait evolution are commonly reported without any estimate of the variance, most likely because the focus is often on the results of model tests and p values rather than in our ability to reliably estimate and interpret the parameters of the model (see discussion in Beaulieu and O’Meara, 2016 on a related issue). Furthermore, model tests such as the likelihood ratio test and the Akaike information criteria (AIC) do not directly incorporate measures of the variance of estimates in their calculations. This is problematic when parameters can be hard to estimate and models are challenged by reduced sample sizes, which is a common issue in phylogenetic comparative methods analyses in general.

The analysis of mouth shape evolution in function of diet in Centrarchidae fishes (Revell and Collar, 2009) is an interesting example of the impact of uncertainty in parameter estimates on our biological conclusions. The likelihood ratio test showed a strong support for a shift in the structure of evolutionary correlation associated with the evolution of piscivory in the *Micropterus* clade. In contrast, the summary statistics computed from the posterior distribution did not show strong evidence for the this scenario. When we contrast the result from the MLE with the posterior distribution (Fig. **??**), we can visualize the origin of the incongruence. The likelihood ratio test focus on the relative fit of the constrained model (one regime) compared with the full model (two regimes) whereas the summary statistics computes whether our knowledge about the model updated by the data reflects a strong signal for a shift between regimes. Furthermore, the same trend can be shown by computing the confidence interval around the MLE estimates using bootstrap simulations (Fig. S9), since there is an important overlap between **R** matrices fitted to each regime.

The results from the test of whether mainland and island anole species differ in the pattern of evolutionary integration among traits are intriguing. There is a change in the pattern of evolutionary correlation between body size and head length associated with the dispersion from the Caribbean islands back to the mainland. However, there is no detectable difference in the pattern of evolution for these traits between the ancestral mainland and island lineages. We estimated a weaker evolutionary correlation between body size and head length in the dispersal mainland group (Figure 5 and Table S1). A tight positive integration between head and body size may allow for the development of extreme head shapes whereas a looser, but still positive, integration observed in ‘dispersal mainland’ species suggests morphological evolution in the clade is spreading faster in the morphospace. It is natural to expect that a shift in the trajectory of evolution of morphological traits associated with ecomorphs would occur, since mainland and island anole species are known to occupy different regions of the morphospace (Pinto et al., 2008; Schaad and Poe, 2010; Moreno-Arias and Calder´on-Espinosa, 2016). Surprisingly, our results suggest that ecomorphs evolved under a constant pattern of evolutionary integration among traits with respect to ancestral mainland lineages. However, changes in the selective regime can be responsible for rapid phenotypic differentiation even in the absence of correspondent shifts in evolutionary correlation among traits (Drake and Klingenberg, 2010; Gerber, 2013). Thus, island and mainland anole lineages are not distinct in their potential to explore the morphospace and ecomorphs might be special in the sense of repetitive radiations, but not due to distinct mode or pace of morphological evolution when compared to their mainland counterparts (also see dicussion by Poe et al., 2018). This explanation has some support by the fact that a few mainland species are morphologically similar to island ecomorphs (Schaad and Poe, 2010). Our results point to the idea that ecomorphs might also have evolved among mainland species, but efforts to understand anole biodiversity, ecology and evolution have been strongly focused on island systems and still relatively very little is known about mainland lineages.

## Conclusion

Most of what we know about the tempo and mode of trait evolution come from studies of individual traits, but evolutionary integration is ubiquitous across the tree of life. Recently we have seen an increase in comparative tools aimed to deal with the challenges posed by high-dimensional traits, such as shape data. However, the discipline is still in need of better models to deal with multiple traits, such as the examples explored in this study. Our framework is aimed primarily on the study of the structure of evolutionary correlation among multiple univariate traits across clades and over time. The implementation of summary statistics make it feasible to extend our approach to be focused on any attribute of the evolutionary rate matrix that might fit the biological predictions of a specific study. Another important advantage of Bayesian MCMC is that proposals can be modified to integrate over different number of regime configurations, distinct models of trait evolution, and even simultaneously estimate parameters for the trait evolution model and the phylogenetic tree. Thus, our implementation lays the groundwork for future advancements towards flexible models to explore multiple facets of the evolution of integration over long time scales using phylogenetic trees. Integration among traits is a broad and yet fundamental topic in evolutionary biology. Understanding the interdependence among traits over the macroevolutionary scale can be key to tie together our knowledge about the genetic basis of traits, development, and adaptive shifts in the strength or direction of evolutionary correlation.

## Acknowledgements

We would like to thank Josef Uyeda, Rosana Zenil-Ferguson, Matthew Pennell, Mason Linscott, Dave Tank, Brian Sidlauskas, and Dean Adams for insightful discussions on the topic and Julien Clavel for comments on an earlier version of this manuscript.

---

**Table S1:**
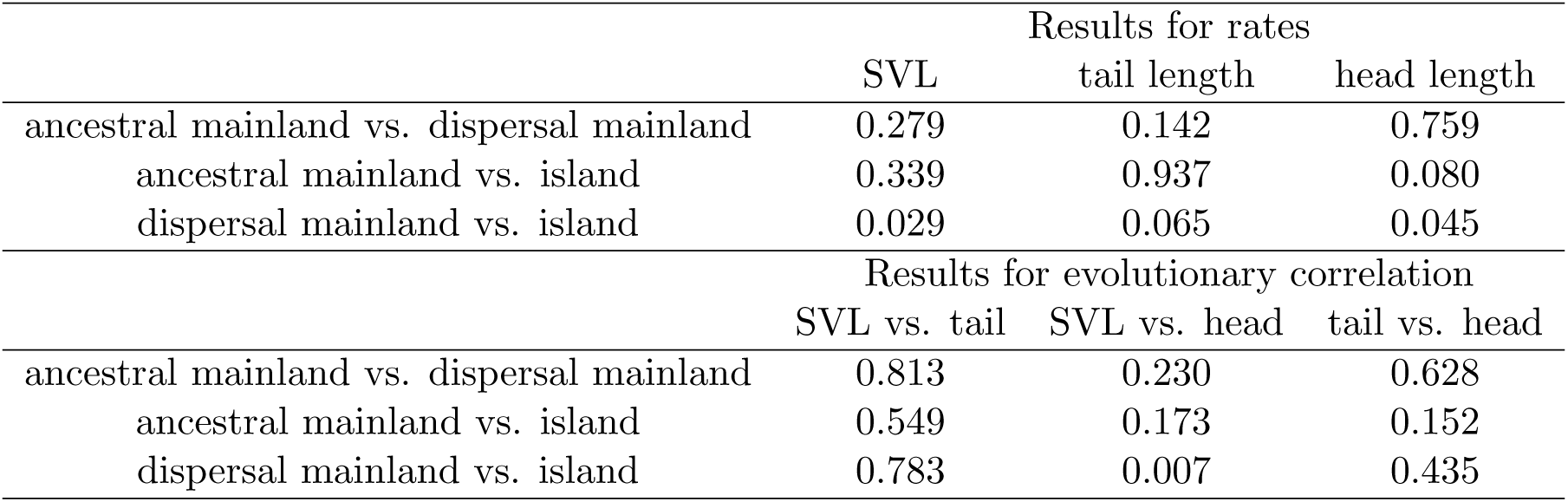
Results from summary statistics quantifying the overlap between the posterior distribution for the pairwise rates of evolution and evolutionary correlations among traits across the three regimes fitted to the time tree of mainland and island anoles.

|  | Results for rates |  |  |
| --- | --- | --- | --- |
|  | SVL | tail length | head length |
| ancestral mainland vs. dispersal mainland | 0.279 | 0.142 | 0.759 |
| ancestral mainland vs. island | 0.339 | 0.937 | 0.080 |
| dispersal mainland vs. island | 0.029 | 0.065 | 0.045 |
|  | Results for evolutionary correlation |  |  |
|  | SVL vs. tail | SVL vs. head | tail vs. head |
| ancestral mainland vs. dispersal mainland | 0.813 | 0.230 | 0.628 |
| ancestral mainland vs. island | 0.549 | 0.173 | 0.152 |
| dispersal mainland vs. island | 0.783 | 0.007 | 0.435 |

**Figure S1:**
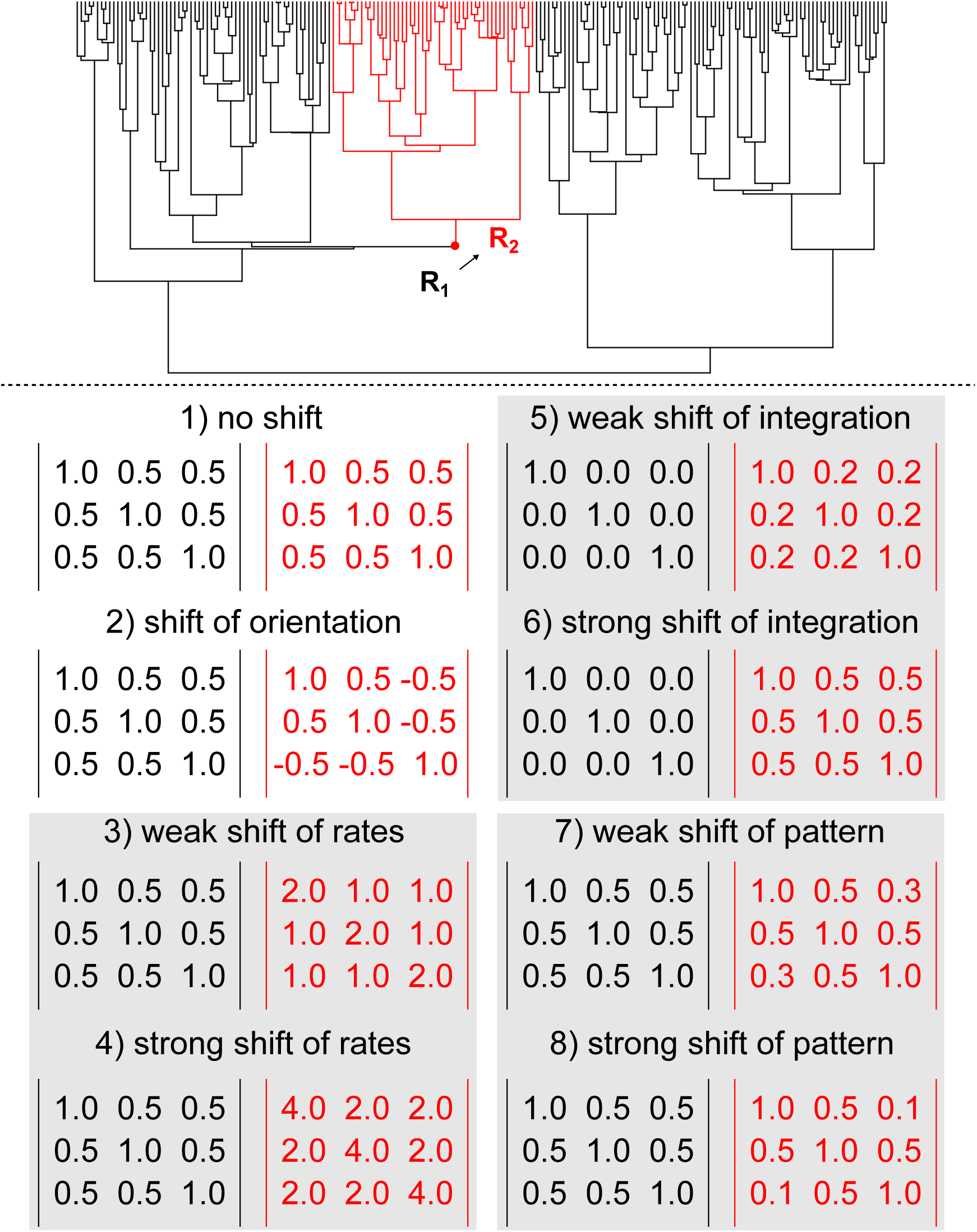
Parameter values used to generate data sets under eight different scenarios of correlated evolution and 3 traits. Phylogenetic tree at the top is an example of a 200 tips tree with a 50 tips focus clade in red. A single shift from the background rate regime (branches and matrices in black) to the focus regime (branches and matrices in red) happen at some point along the steam branch leading to the focus clade. Matrices show the parameter values for the multivariate Brownian motion process used to generate data sets for each of the eight distinct scenarios. Grey squares group simulation treatments under the same type of evolutionary correlation. All traits were simulated with root value equal to 10.

**Figure S2:**
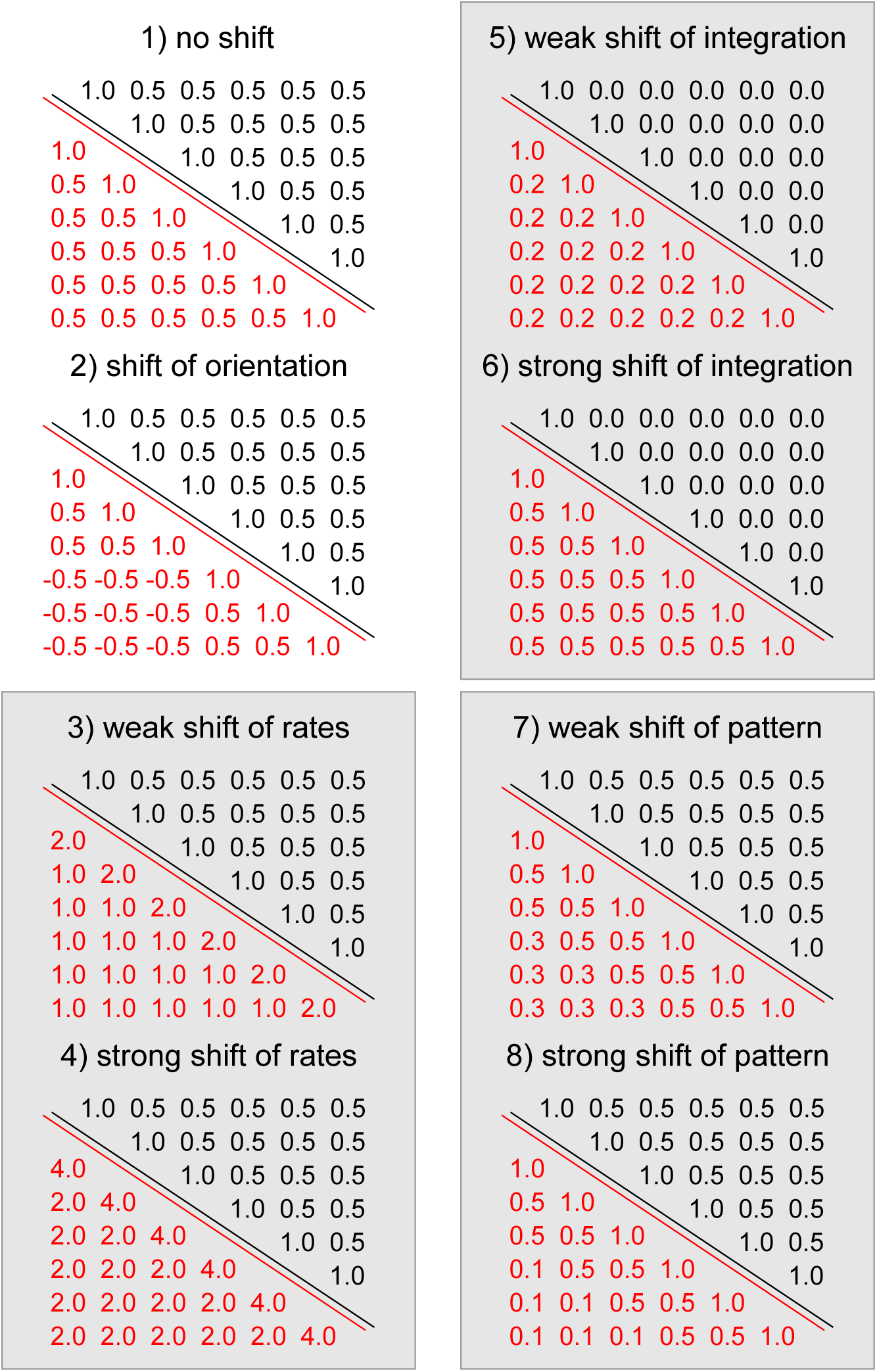
Parameter values used to generate data sets under eight different scenarios of correlated evolution and 6 traits. Upper and lower triangular matrices show the parameter values for the multivariate Brownian motion process used to generate data sets for each of the eight distinct scenarios. Upper triangular matrices in black were used to generate data for background regime whereas lower triangular matrices in red were used in the focus clades (see Figure S1). Grey squares group simulation treatments under the same type of evolutionary correlation. All traits were simulated with root value equal to 10.

**Figure S3:**
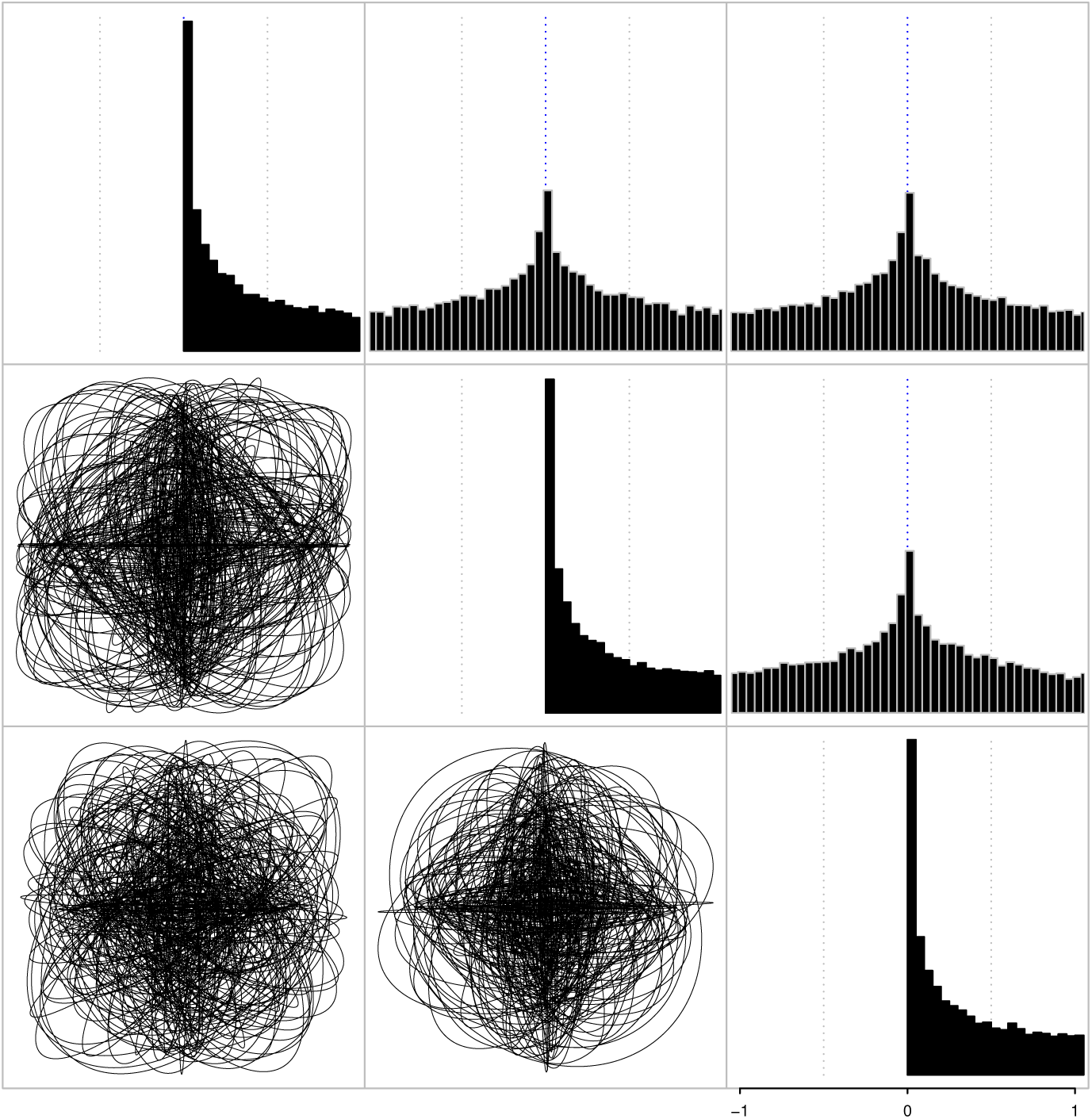
Prior distribution for the evolutionary rate matrix (**R**) used for all analyses. Plate shows samples in the interval between -1 and 1 from the prior for a model with three traits. Diagonal plots represent the prior for evolutionary rates (variances) for each trait, upper-diagonal plots show pairwise evolutionary covariation (covariances), and lower-diagonal are samples from the posterior distribution of ellipses showing the 95% confidence interval of each bivariate distribution.

**Figure S4:**
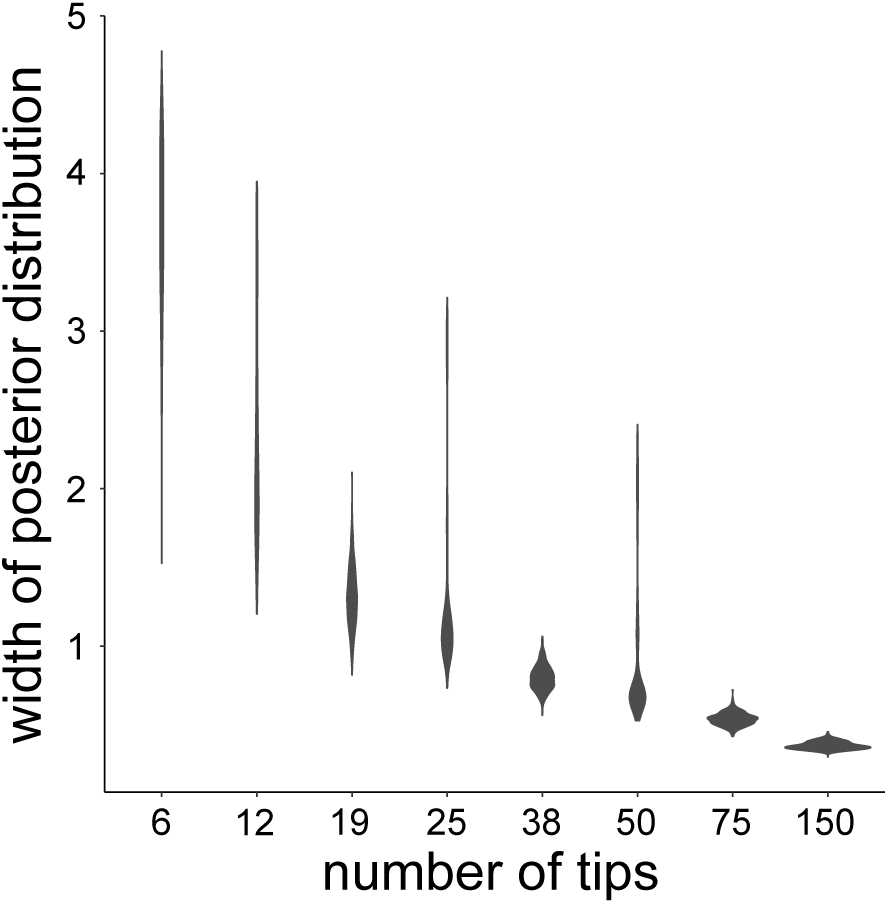
Uncertainty in the posterior distribution of parameter estimates in function of the number of species in the phylogeny. The y axis shows the average width of the posterior distributions among all parameters of the model (rates, correlations, and root values) for each replicate across all scenarios (total of 800 per phylogeny size).

**Figure S5:**
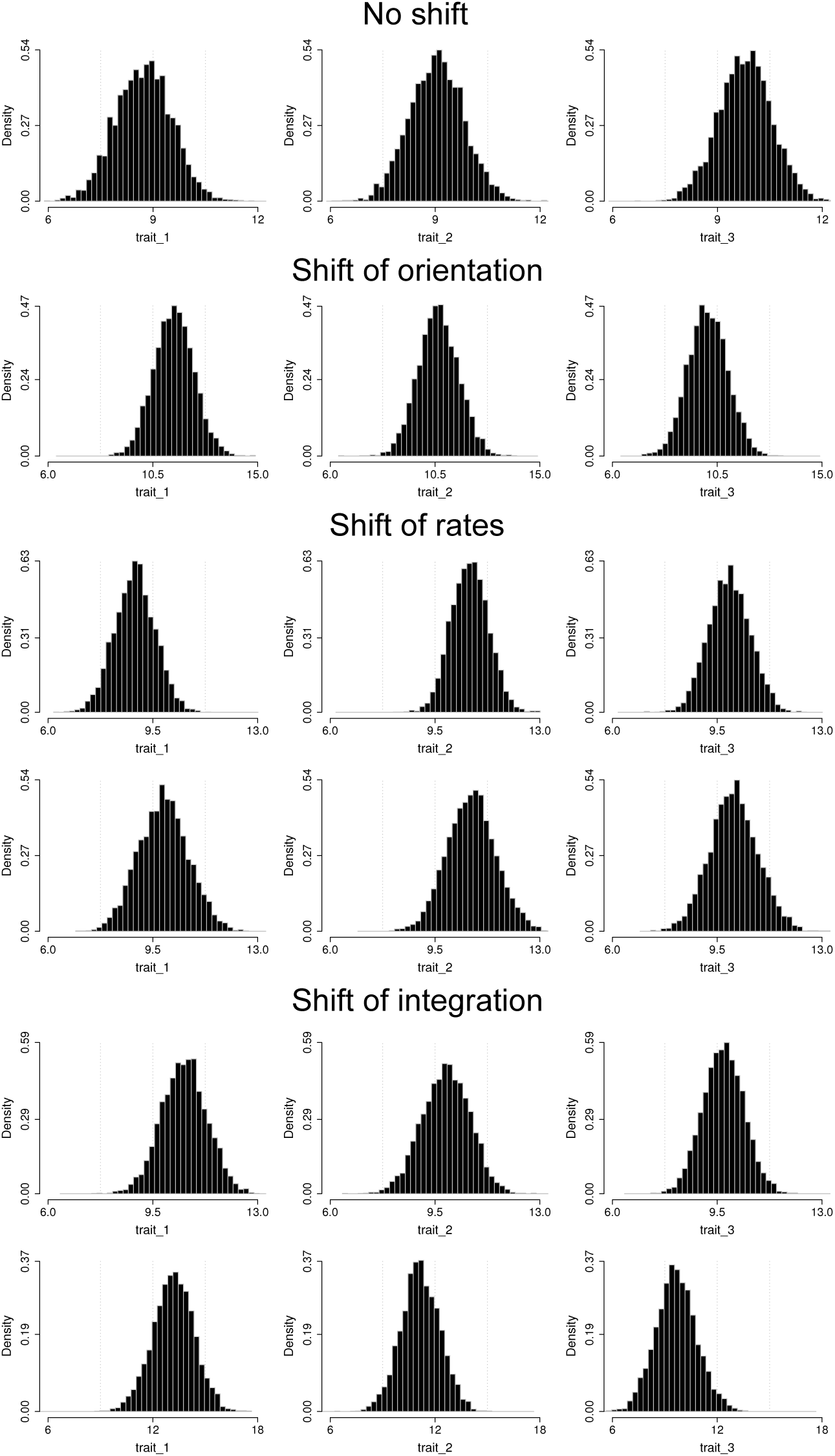
Example of posterior distribution of root values for six of the eight simulation treatments with three traits each. Simulation treatments are the same as showed on Figure 2 and 3. Top and bottom plots for ‘Shift of rates’ and ‘Shift of integration’ treatments correspond to the weak and strong shifts of the same treatments, respectively (see Figures 2 and 3). The true value for the ancestral state of each trait in all simulations was equal to 10. Note that results are congruent with when the root prior is an (improper) uniform prior as shown on Figure S6.

**Figure S6:**
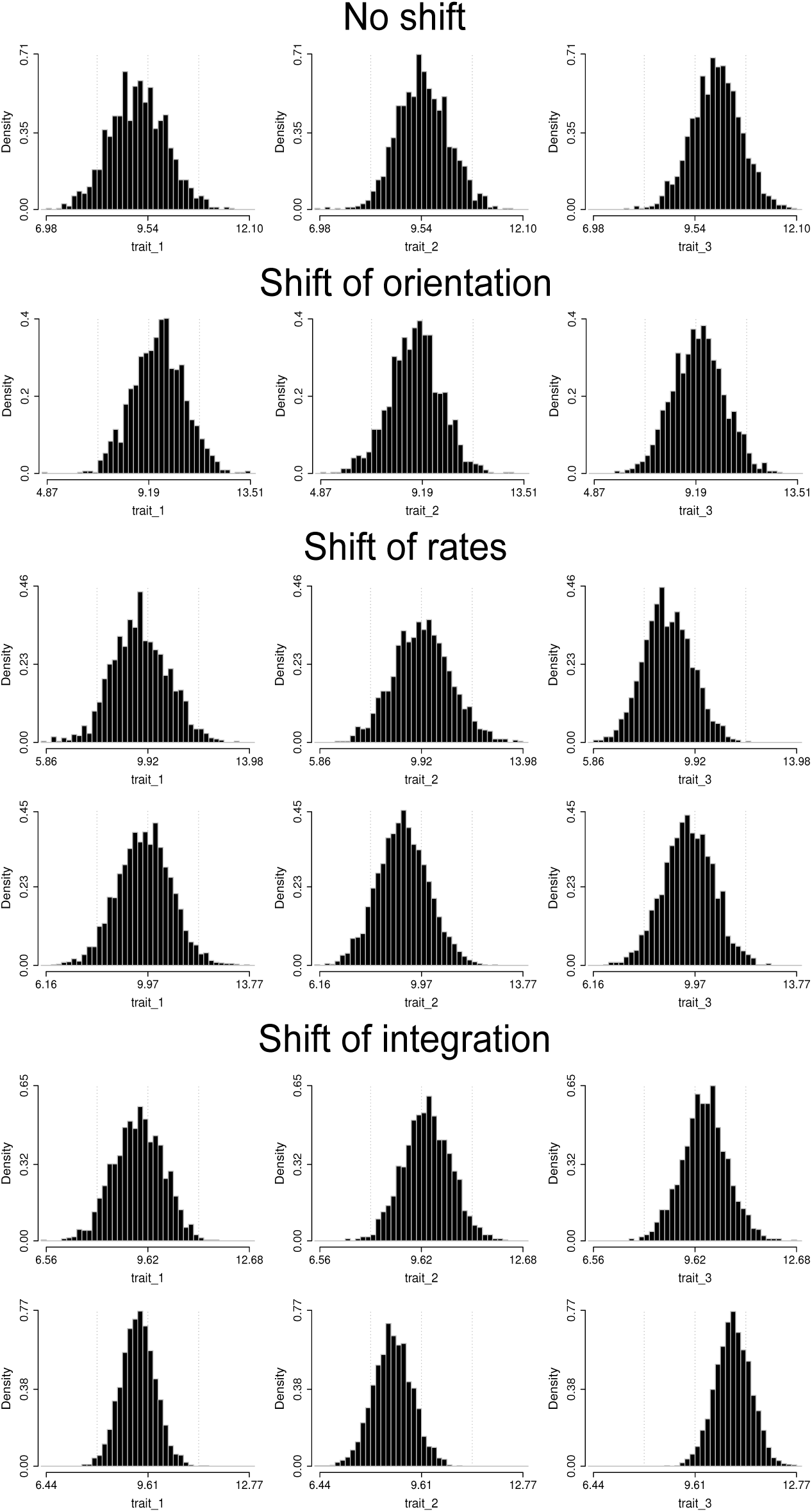
Example of posterior distribution of root values for six of the eight simulation treatments with three traits each using an (improper) uniform prior on the root. Simulation treatments are the same as showed on Figure 2 and 3. Top and bottom plots for ‘Shift of rates’ and ‘Shift of integration’ treatments correspond to the weak and strong shifts of the same treatments, respectively (see Figures 2 and 3). The true value for the ancestral state of each trait in all simulations was equal to 10. Note that results are congruent with when the root prior is normally distributed with mean and standard deviation informed by the tip data as shown on Figure S5.

**Figure S7:**
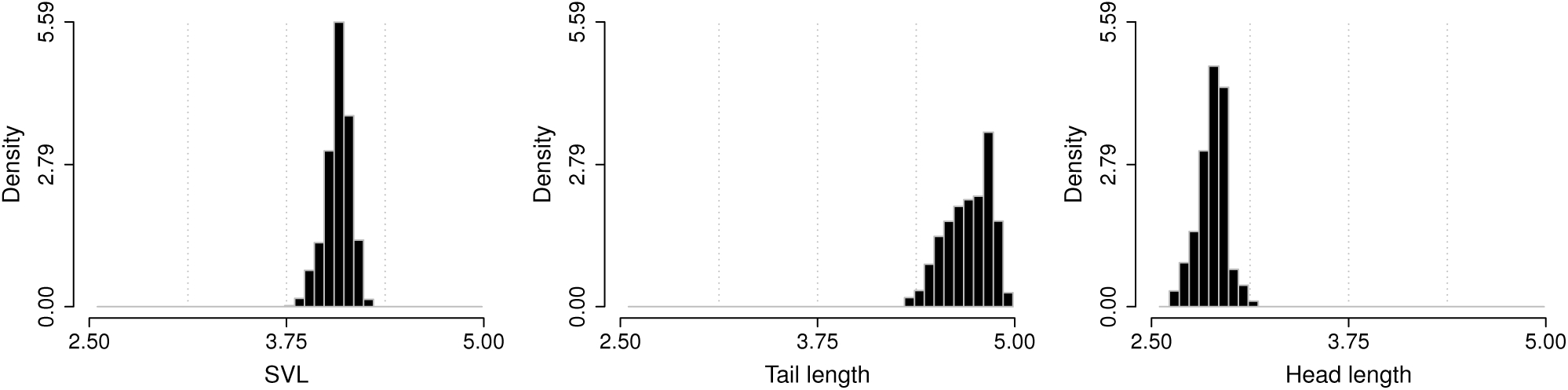
Posterior distribution of root values for anoles.

**Figure S8:**
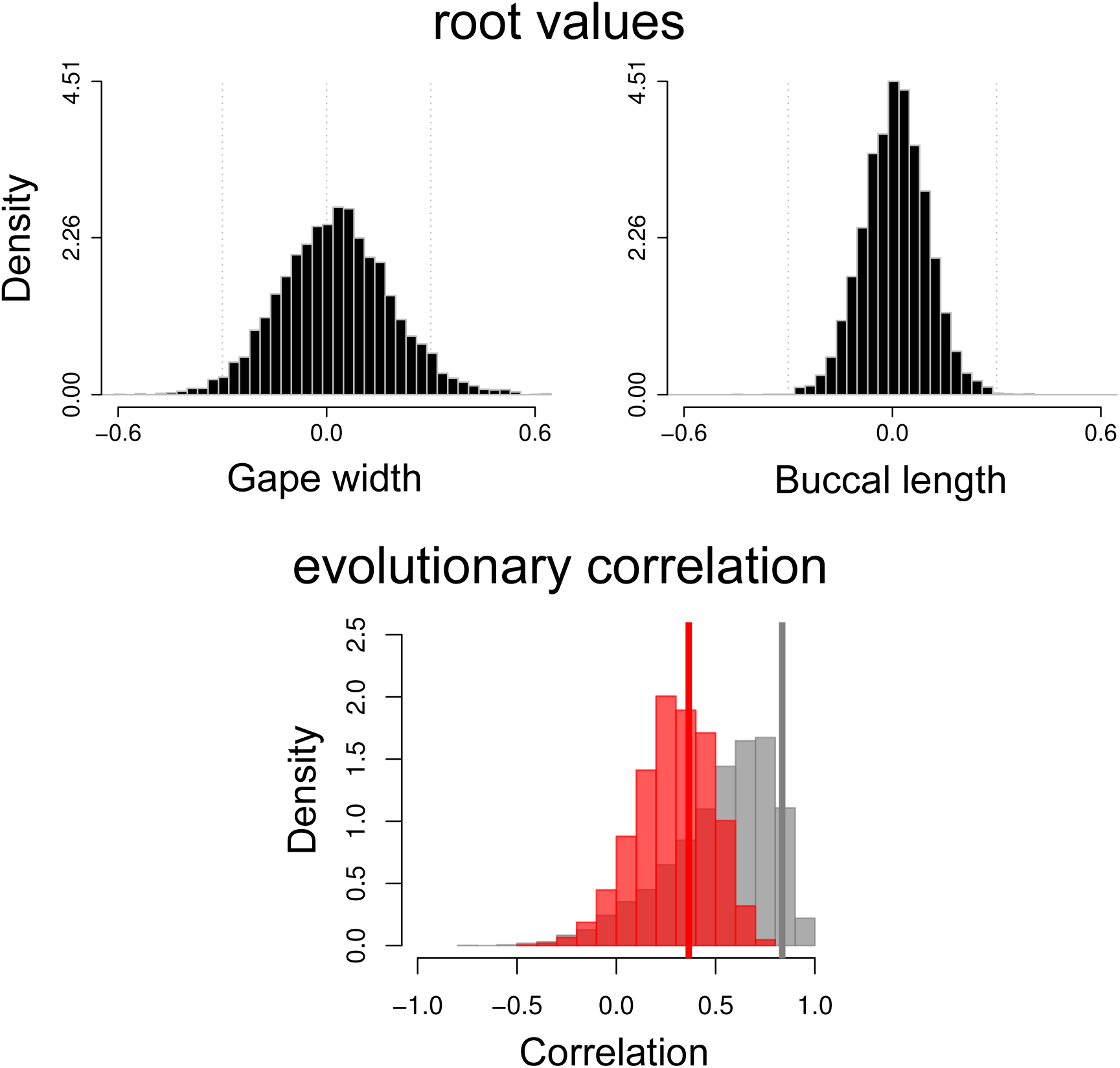
Posterior distribution of root values and evolutionary correlation fitted to the Centrarchidae fishes. Evolutionary correlation between gape width and buccal length estimated for the background group showed in gray and for the *Micropterus* clade in red. Vertical lines matching regime colors show the point estimate for the evolutionary correlation using maximum likelihood. See Figure 6 for the posterior distribution of evolutionary rate matrices for each regime and the phylogeny used for the analysis.

**Figure S9:**
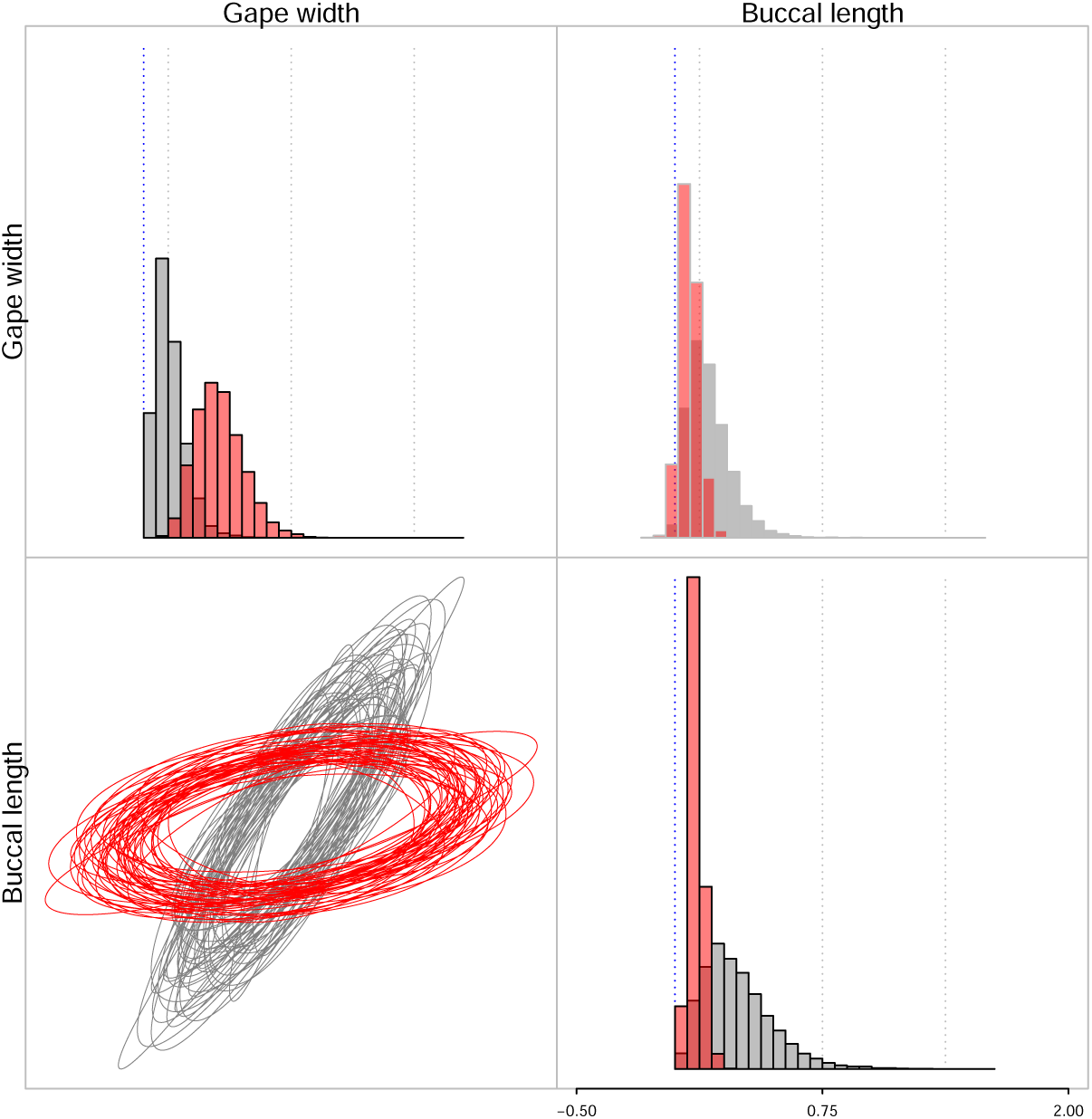
Distribution of parameter estimates from 10000 parametric bootstrap simulations for the two regimes model. Background group showed in gray and *Micropterus* clade showed in red. Diagonal plots show evolutionary rates (variances) for each trait, upper-diagonal plot show pairwise evolutionary covariation (covariances), and lower-diagonal plot shows samples from the posterior distribution of ellipses showing the 95% confidence interval of each bivariate distribution.

## Supplementary material for

### The pruning algorithm used to calculate the likelihood of multiple R matrices fitted to the tree

Here we describe in details the pruning algorithm (Felsenstein, 1973, 1985) applied to calculate the log-likelihood of a multivariate Brownian motion model in which the rates vary throughout the branches of the tree. A script for the R programming language with a commented implementation and example of the algorithm was provided as online supplementary material file in the Dryad repository (see link in main text).

The pruning algorithm explores the property that trait changes in each of the branches of a phylogenetic tree can be modelled independently and applies a multivariate normal density to compute the likelihood of evolutionary changes through time assuming a Brownian motion model (Felsenstein, 2004; Freckleton, 2012) of continuous character evolution. When multiple rate regimes are fitted to a phylogeny, the likelihood is often computed by scaling each branch of the tree by its corresponding rate of evolution and then applying the pruning algorithm with evolutionary rate parameter equal to 1 (e.g., Eastman et al., 2011). By doing this we can implement the pruning algorithm for a single rate Brownian motion even though the model has multiple evolutionary rates. However, this procedure is not applicable to the multivariate case, since the product of the length of a branch and the BM rate is a matrix which makes scaling the branches of the phylogeny impossible. As a result, we need to extend the pruning algorithm to the case of multiple evolutionary rates for a multivariate Brownian motion model. We derived the pruning algorithm for multiple rate regimes by following the same procedures described by Felsenstein (1973, 2004), but assuming that all rates are multivariate, that rates can be different on each branch and that branches can have multiple rate regimes (after Revell and Collar, 2009). This algorithm completely avoids the calculation of the matrix inverse and the determinant of the phylogenetic covariance matrix (**C**) or the Kronecker product between **R** and **C** matrices. However, the inverse of the **R** matrix, which will have dimensions equal to the number of traits in the data set, is still required.

In this extension of the algorithm, each branch of the phylogeny can be assigned to one or more evolutionary rate matrix (**R**) regimes and the sum of the portions of the branch assigned to each regime has to be equal to the total length of that branch (Revell and Collar, 2009). We demonstrate that the algorithm yields the same likelihood as in Felsenstein (1973) and Freckleton (2012) by showing that all calculations converge when a single regime is fitted to the tree. The pruning algorithm works by visiting the tips and going down node by node. At each step the contrast between two tips is computed and a new “phenotype” value replaces the two original tips, becoming the new tip. The likelihood of the contrast is calculated and we move to the next contrast until we reach the root node. From here on we will refer to the following phylogenetic tree as an example:

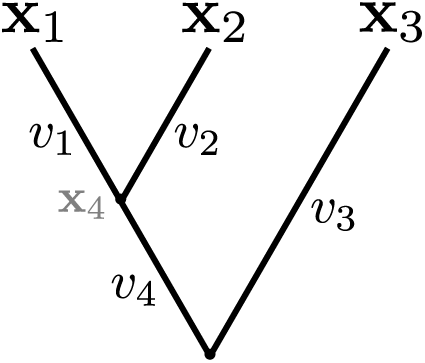

Where **x**_*i*_is a vector with *r* trait values for tip *i* and *v*_*i*_ is the branch length leading to tip or node *i*. We will refer to the node representing the common ancestor of tips 1 and 2 as the node 4 and the node representing the common ancestor of all tips as the root node. The method works as following:

1. **Calculate the contrast.** Choose a pair of tips *i* and *j* with a unique and exclusive common ancestor *k*. In our example, the selected species are 1 and 2. Compute the contrast **u**_*ij*_= **x**_*i*_ − **x**_*j*_.
2. **Compute the log-likelihood.** Use the vector of contrasts (**u**_*ij*_), the number of traits in the data (*r*), the branch lengths (*v*_*i*_ and *v*_*j*_), and the length of the branches assigned to each of the *k* evolutionary rate matrix regimes (Revell and Collar, 2009) to compute the log-likelihood:

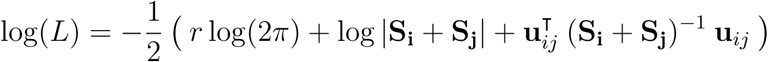

where

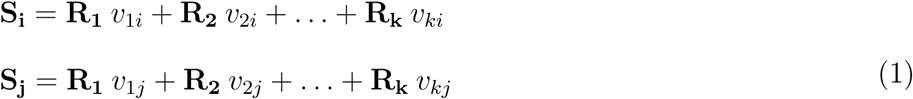

and

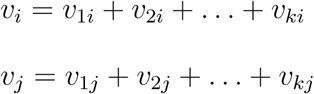 If we assume a single evolutionary rate matrix is fitted to the whole tree, equation 1 reduces to equation 10 in Freckleton (2012):
Let

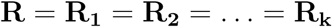

then

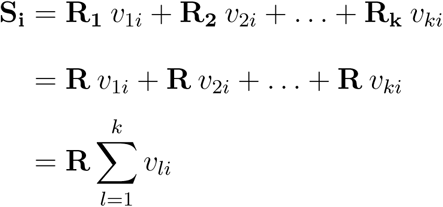 We know, from equation 1, that the sum of the portions of the branch length assigned to each regime is equal to the total length of the branch. Then:

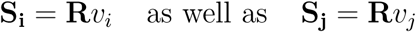

and

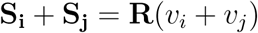 Substituting into equation 1, we have:

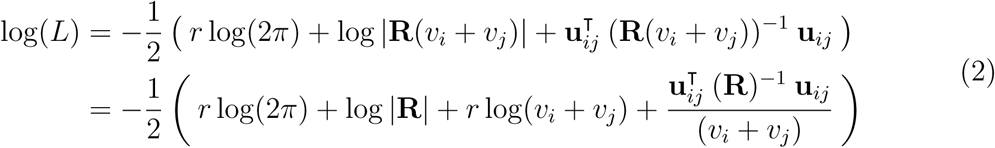 Which is the same as equation 10 in Freckleton (2012)^1^.
3. **Calculate the new phenotype vector x**_*n*_**for the node** *n*. This quantity is originally calculated as the weighted average of the vector of species means for species *i* and *j* with weights equal to the length of the branches *v*_*i*_ and *v*_*j*_. For the case of a single trait, **x**_1_and **x**_1*j*_, we would have:

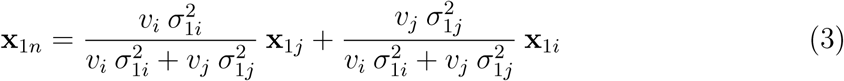 When 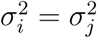, equation 3 becomes equivalent to equation 7 in Felsenstein (1973) and the rates of each branch can be omitted. However, here we assume that rates can be different in every branch, that the evolutionary covariance among traits are non-zero and that more than one rate regime can be assigned to the same branch. As a result, the rates need to be represented as variance-covariance matrices for the regimes 1 to *k* (**R_1_**, **R_2_**, …, **R_k_**) and the sum of the product between the portions of each branch and their rate regimes is given by the matrices **S_i_** and **S_j_** (see equation 1). By expanding equation 3, we have:

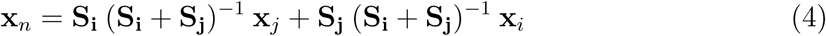 In the first step of our example, we calculated the phenotype value for the node 4 (**x**_4_). Then, we prune the tips 1 and 2 from the tree, leaving only the tip 3 and the new tip 4 with vector of trait values **x**_4_. The next contrast will be calculated between **x**_4_ and **x**_3_, leading to the phenotype value for the root node of the tree.
4. **Compute the variance of x**_*n*_. After computing the vector of trait values for the node *n*, we need to calculate the variance associated with the uncertainty in the estimation of **x**_*n*_. This uncertainty is added to the variance of the branch immediately bellow the node *n*. For a single trait and we would have:

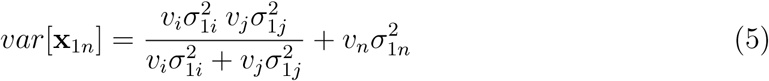 Where *m, …, n* are the indexes for the branches that connect the root to the node *n* of the tree. Again, when a single rate regime is fitted to the tree, equation 5 is equivalent to equation 9 in Felsenstein (1973). For the multivariate case, this quantity becomes a variance-covariance matrix which is added to **S_n_** (the branch length below the node *n* multiplied by the rate regimes; see equation 2) and can be calculated as:

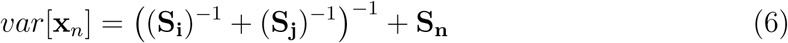 The equivalence between equations 5 and 6 can be easily verified by checking the computation of the harmonic mean of matrices. For two scalar quantities the harmonic mean is given by 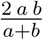 whereas for matrices we have 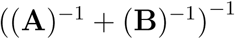.
5. **Repeat.** Steps 1 to 4 are repeated until only two tips remain.
6. **Root value.** The ancestral node of the two remaining tips is the root node of the phylogeny. We can compute the phenotype value for the root node (**x_root_**) using step 3. The variance for this estimate is computed as:

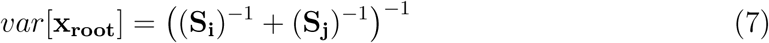 The log-likelihood for the root is computed as:

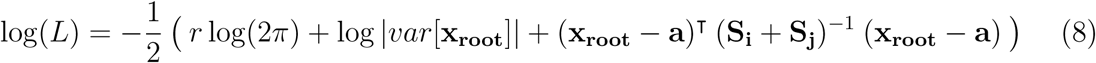 Where **a** is a vector with *r* root values for the tree. Thus, (**x_root_** *−* **a**) is the difference between the estimated value for the root and the parameter for the multivariate Brownian motion model (**a**).
7. **Compute the final log-likelihood.** The final log-likelihood conditioned on the phylogenetic tree, rate regime, trait data and root value is computed as the sum of all partial (node-by-node) log-likelihoods computed in step 2 and the log-likelihood for the root node computed in step 6.

## Footnotes

1 Note that the published equation in Freckleton (2012) has a printing error. The corrected form is 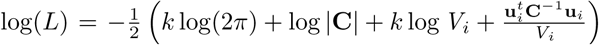. The correct form can be appreciated in the function ‘clikGeneral’ on line 393 of the Supporting Information file MEE3 220 sm demo.R available online (Freckleton, 2012).

